# BORC segregates synaptic vesicle and lysosomal proteins through motors UNC-104/KIF1A and UNC-116/KIF5

**DOI:** 10.1101/2024.12.31.630845

**Authors:** Amal Mathew, Sohan Seal, Aditee Dandekar, Badal Singh Chauhan, Sruthi Sivadasan, Michael L Nonet, Sandhya P. Koushika

**Affiliations:** Department of Biological Sciences, Tata Institute of Fundamental Research, Mumbai, 400005, India; Department of Neuroscience, Washington University School of Medicine, St Louis, MO, 63110, U.S.A

**Keywords:** : BORC, SAM-4, Myrlysin, ARL8, LRK-1, LRRK2, AP3, APB-3, synaptic vesicle biogenesis, lysosomal proteins

## Abstract

While synaptic vesicle proteins (SVPs) and lysosomal proteins can be present together in axons, the origin of these compartments is unclear. These SV-lysosomes are however thought to segregate further to SVPs and lysosomal proteins. In this study, we identify genes and characterize a genetic pathway involved in the segregation of SVPs and lysosomal proteins in the neuronal cell body. We identify a novel role for BORC (BLOC-1-related complex) in segregating SVPs and lysosomal proteins in the cell body. BORC subunit SAM-4/Myrlysin acts through ARL-8 and Kinesin motor proteins UNC-116/KIF5 and UNC-104/KIF1A in segregating SVPs and lysosomal proteins. Additionally, we also show that LRK-1/LRRK2 and APB-3/AP-3 (β3), involved in pre-SV biogenesis, regulate the segregation of SVPs and lysosomal proteins in the neuronal cell body. LRK-1 recruits SAM-4 that in turn governs the localisation of APB-3 suggesting a hierarchical pathway of LRK-1-SAM-4-APB-3 for the segregation of SVPs and lysosomal proteins. Additionally, we also observe that the size of lysosomal protein-containing compartments (LPCCs) is smaller in SAM-4 and LRK-1. This size regulation depends on UNC-116. Together, we show that BORC recruited by LRK-1 and in turn via motors and along with AP-3 mediates the segregation of SVPs and lysosomal proteins in the neuronal cell body.

## Introduction

Neuronal function requires sorting of distinct cargoes to their appropriate destinations [1,2]. Secretory cargo can be sorted at the endoplasmic reticulum (ER), at the trans-Golgi network (TGN) or at post-Golgi compartments [3,4]. Synaptic vesicles are a major cargo in neurons enriched at axons and synapses. The precursors of synaptic vesicles (pre-SVs) are made in the cell body and are transported to synapses by the Kinesin-3 motor protein KIF1A/UNC-104 [5,6].

Several early steps in synaptic vesicle precursor (pre-SV) biogenesis have been identified, namely the exclusion of Golgi proteins, inclusion of synaptic vesicle proteins, and regulation of pre-SV size [7]. These processes are regulated by UNC-16/JIP3 and LRK-1/LRRK2 via the clathrin adaptor protein complexes AP-3 and AP-1 respectively governing in axons pre-SV membrane protein composition and compartment size [7,8]. Additionally, by quantifying the cotransport of SVPs and lysosomal proteins in axons of *C. elegans*, we identified LRK- 1/LRRK2-mediated segregation of SVPs and lysosomal proteins via AP-3 as a critical step in pre-SV biogenesis [8]. While lysosomal and SVPs do not co-transport in axons of rat hippocampal neurons, and their co-transport is debated in mouse hippocampal neurons, human iPSC-derived, *Drosophila*, and *C. elegans* neurons co-transport lysosomal and synaptic vesicle proteins [8–11]. These axonal SV-lysosomes may originate from the cell body or they could arise from retrogradely moving lysosomes fusing with pre-SVs [8,12–14]. While SVP and lysosomal proteins are thought to be separated from common compartments in the cell body, we do not understand events in the cell body that lead to this separation [8,12,14]. Additionally, the transport of SVPs and lysosomal proteins in distinct carriers in some vertebrate neurons could arise as a consequence of segregation events in the cell body. The current studies in the field warrant the assessment of SVP and lysosomal trafficking in the cell body, a hub for the biogenesis of both pre-SVs and lysosomes.

Genes that could regulate the trafficking of both SVPs and lysosomal proteins might be involved in the segregation of SVPs and lysosomal proteins in the cell body. One such complex, BORC (BLOC-1 related complex), a multi-subunit complex, was initially identified for its role in lysosomal positioning in non-neuronal and neuronal cells [15]. In mouse hippocampal neurons, BORC is dispensable for SVP transport, but in human iPSCs, BORC mutants rescue ARL-8 KO synaptic vesicle transport defects by increasing KIF1A recruitment via elevated PI(3,5)P_2_ levels [10,11]. Interestingly, in *C. elegans* neurons, SVP distribution partially depends on SAM- 4/Myrlysin, a BORC subunit [16,17]. These findings suggest that BORC regulates axonal SVP distribution and lysosomal distribution differently across model systems. BORC acts through the GTPase ARL-8 and motor proteins Kinesin-3 and Kinesin-1 to regulate the distribution of SVPs and lysosomal proteins [15–19]. While Kinesin-3 regulates the distribution of both lysosomal and synaptic vesicle proteins in mouse hippocampal neurons and human iPSCs, Kinesin-1/UNC-116 controls lysosomal protein distribution [9–11,19].

Since segregating lysosomal and synaptic vesicle proteins may be a key step in pre-SV biogenesis, we examined the role of BORC, ARL-8, and ARL-8 dependent motor proteins in this process in the neuronal cell body. We show that the segregation of SVP and lysosomal proteins in the neuronal cell body is reduced in BORC mutants. Further, BORC acts through the GTPase ARL-8 and ARL-8 dependent motor proteins, UNC-104/KIF1A and UNC-116/KIF5, in segregating SVP and lysosomal proteins. We also show that SAM-4 through ARL-8 and UNC- 116 also regulate the size of lysosomal protein-containing compartments (LPCCs). SAM-4 localisation is regulated by LRK-1 and SAM-4 in turn regulates the localisation of APB-3 placing SAM-4 below LRK-1 and ahead of AP-3 in the hierarchical pathway. Together, our study uncovers a novel role for BORC acting through the motor proteins UNC-104 and UNC- 116 in segregating SVPs and lysosomal proteins and regulating the size of lysosomal compartments and enabling exit of LPCCs into the axon.

## Results

### Segregation of synaptic vesicle proteins and lysosomal proteins in the neuronal cell body is dependent on BORC but not BLOC-1

BORC is a multisubunit complex containing the following subunits: *blos-1/*BLOS1, *blos- 2/*BLOS2, *snpn-1/*Snapin, *sam-4/*Myrlysin, *blos-7/*Lyspersin, *blos-9/*MEF2BNB, *kxd-1/*KXD1 and *blos-8/*Diaskedin [15,17]. Three of these subunits (*blos-1/*BLOS1, *blos-2/*BLOS2, and *snpn- 1/*Snapin) are shared with the BLOC-1 (Biogenesis of lysosome-related organelle complex 1) complex [15]. To investigate the role of BORC and BLOC-1 in segregating SVPs and lysosomal proteins we investigated the overlap of synaptic vesicle proteins and lysosomal proteins in the cell body of the posterior lateral mechanosensory (PLM) neuron of *C*. *elegans* (S1A).

We examined the overlap of different pairs of synaptic vesicle and lysosomal proteins in the cell body of PLM in the mutants of 2 subunits of BORC, *sam-4*/Myrlysin and *snpn-1*/Snapin-1. We examined the overlap of two transmembrane proteins, lysosomal cystine transporter Cystinosin (CTNS-1) and synaptic vesicle protein synaptogyrin-1 (SNG-1), in two different null alleles of *sam-4(js415)* and *sam-4(tm2838)* both represented as *sam-4(0)* [16]. We observed an increase in the overlap of SNG-1 tagged with mNeonGreen and CTNS-1 tagged with mCherry throughout the volume of the cell as indicated by an increase in the Pearson’s and Mander’s coefficients (Fig 1A, 1B, 1C; S1A, S1B, S1C, Movie 1 and 2). In *snpn-1(tm1892)* mutants, we observed a similar increase in the overlap of SVPs and lysosomal proteins (Movie 3 and 4). We also observed an increase in the overlap of the synaptic vesicle protein synaptobrevin (SNB-1) and lysosomal lysine/arginine transporter (LAAT-1) through out the volume of the cell in *sam-4(0)* indicating that the segregation defects are independent of the SVP and lysosomal protein pair (S1E, S1F, and S1G, Movie 5 and 6). By contrast, we did not detect any increase in the overlap of SNG-1 and CTNS-1 in BLOC-1 specific subunit mutant, *dsbn-1*, suggesting that the increase in overlap of SVPs and lysosomal proteins is BORC dependent and BLOC-1 independent (Fig 1A, 1B and IC).

**Figure 1:**
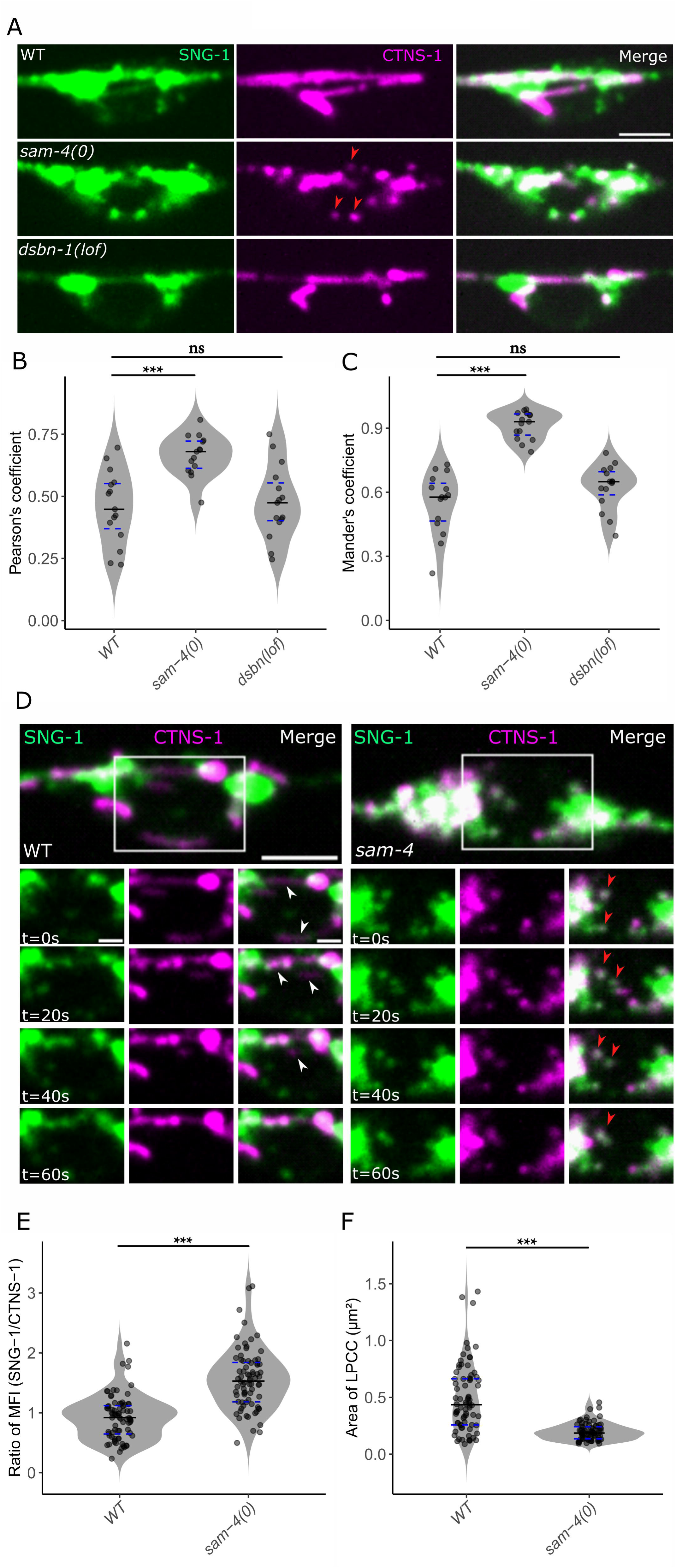
BORC segregates SVPs and lysosomal proteins and regulates the size of CTNS-1 compartments A) Representative images of wild type, *sam-4(js415)* and *dsbn-1(tm6580)* showing the overlap of SNG-1::mNeonGreen (Green) and CTNS-1::mCherry (Magenta) in the PLM cell body. Red arrowheads indicate fragmented CTNS-1 compartments. Scale bar = 2.5 µm. B) Quantification of the overlap of SNG-1 and CTNS-1 by Pearson’s Coefficient in wild type, *sam-4(js415)* and *dsbn-1(tm6580)*. Number of animals = 15, Statistical test : Unpaired Student t-test. C) Quantification of the overlap of SNG-1 and CTNS-1 by Mander’s Coefficient in wild type, *sam-4(js415)* and *dsbn-1(tm6580)*. Number of animals = 15, Statistical test : Unpaired Student t-test. D) Representative images of wild type and *sam-4(js415)* showing the overlap of SNG- 1::mNeonGreen (Green) and CTNS-1::mCherry (Magenta) in dynamic compartments in the PLM cell body at time points t=0s to t=60s. Left set of panels corresponds to wild type and right set of panels corresponds to *sam-4(js415)*. White arrowheads indicate compartments showing reduced SNG-1 with CTNS-1 while red arrowheads indicate compartments showing fragmented CTNS-1 with higher levels of SNG-1. Scale bar = 2.5 µm (large panel with ROI), 1 µm (time series panels). E) Quantification of the ratio of Mean Fluorescence Intensity (SNG-1/CTNS-1) in dynamic CTNS-1 compartments in wild type and *sam-4(js415)*. Number of animals ≥ 11, Number of compartments ≥ 77, Statistical test : Wilcoxon rank-sum test. F) Quantification of the area of dynamic CTNS-1 compartments in wild type and *sam-4(js415)*. Number of animals ≥ 11, Number of compartments ≥ 77, Statistical test : Wilcoxon rank-sum test. All data plotted as violin plots showing individual data points marked along with the median (solid line), 25th and 75th percentile marked (dashed lines). * p <0.05, ** p <0.01; *** p <0.001, ns - not significant.

We also examined the overlap of SNG-1 and CTNS-1 in dynamic CTNS-1 compartments by measuring the ratio of mean fluorescence intensity (MFI) of SNG-1 and CTNS-1. We observed an increase in the ratio of MFI (SNG-1/CTNS-1) of dynamic CTNS-1 compartments in the cell body of *sam-4(0)* mutants (Fig 1D, 1E; Movie 7 and 8). Increased ratio of MFI (SNG-1/CTNS-1) in *sam-4* mutants also indicates that SVPs and lysosomal proteins move together highlighting that it is not by chance that these proteins are co-localised. While wild type contains tubular CTNS-1 compartments/LPCCs, the compartments in *sam-4(0)* are fragmented and moving CTNS-1 compartments are smaller in size compared to wild type (Fig 1D and 1F). These observations are consistent with the known roles of BORC in regulating late endosomal size in Hela cells [20].

Together, these experiments demonstrate that BORC regulates the segregation of SVPs and lysosomal proteins in the neuronal cell body and the size of CTNS-1 compartments/LPCCs.

### BORC acts through ARL-8 in segregating synaptic vesicle proteins and lysosomal proteins and regulating the size of CTNS-1 compartments

ARL-8, a GTPase, facilitates the trafficking of endolysosomal and synaptic vesicle proteins by linking these organelles to motor proteins [17,18,21–24]. BORC, an upstream regulator of ARL- 8, governs its localisation to lysosomes and synaptic vesicles [15,17]. To determine whether BORC acts through ARL-8 in segregating SVPs and lysosomal proteins, we asked: i) Do *arl- 8(tm2388)* [*arl-8(lof)*] animals show segregation and fragmentation defects observed in *sam-4* mutants? ii) Does the gain-of-function (gof) allele, *arl-8(jpn1)*, encoding a GTP-locked constitutively active ARL-8 improve segregation of SVPs and lysosomal proteins in *sam-4(0)* [25]?

We examined the overlap of SNG-1::mNeonGreen and CTNS-1::mCherry in the PLM neurons of *arl-8(lof)* animals. Similar to *sam-4(0)*, *arl-8(lof)* mutants displayed increased overlap of SNG-1 and CTNS-1 as observed by an increased Pearson’s and Mander’s colocalisation coefficients, suggesting reduced segregation of SVPs and lysosomal proteins (S3A, S3B, Movie 9). The mutants also showed an elevated ratio of MFI (SNG-1/CTNS-1) in moving LPCCs, reflecting increased SVP presence in CTNS-1 compartments (Fig 2A and 2B; Movie 10).

**Figure 2:**
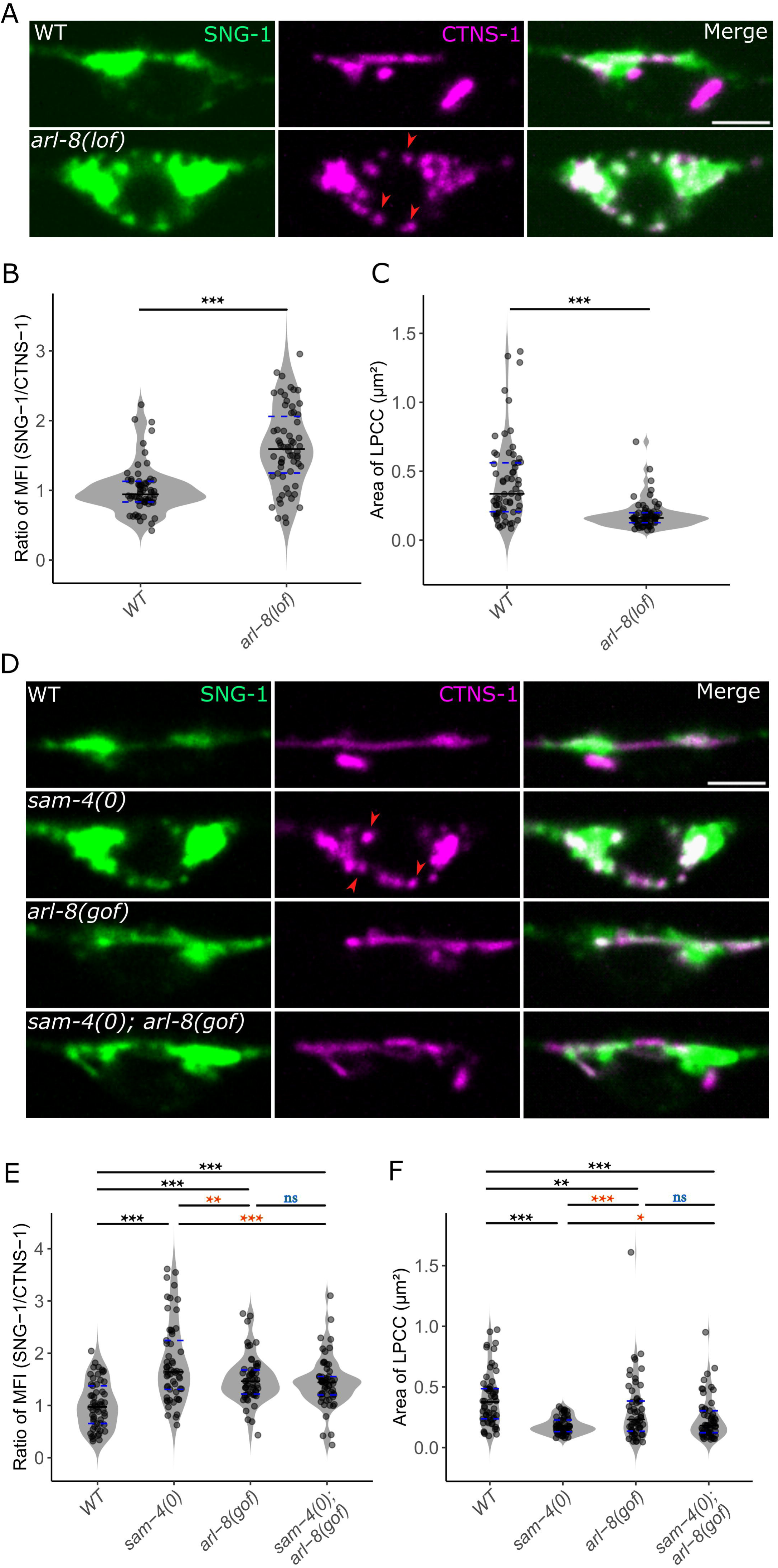
**BORC acts through ARL-8 in segregating SVPs and lysosomal proteins and regulating the size of CTNS-1 compartments** A) Representative images of wild type and *arl-8(tm2388)* showing the overlap of SNG- 1::mNeonGreen (Green) and CTNS-1::mCherry (Magenta) in the PLM cell body. Red arrowheads indicate fragmented CTNS-1 compartments. Scale bar = 2.5 µm B) Quantification of the ratio of Mean Fluorescence Intensity (SNG-1/CTNS-1) in dynamic CTNS-1 compartments in wild type and *arl-8(tm2388)*. Number of animals ≥ 11, Number of compartments ≥ 62, Statistical test : Wilcoxon rank-sum test. C) Quantification of the area of dynamic CTNS-1 compartments in wild type and *arl-8(tm2388)*. Number of animals ≥ 11, Number of compartments ≥ 62, Statistical test : Wilcoxon rank-sum test. D) Representative images of wild type, *sam-4(js415), arl-8(jpn1), sam-4(js415); arl-8(jpn1)* showing the overlap of SNG-1::mNeonGreen (Green) and CTNS-1::mCherry (Magenta) in the PLM cell body. Red arrowheads indicate fragmented CTNS-1 compartments. Scale bar = 2.5 µm E) Quantification of the ratio of Mean Fluorescence Intensity (SNG-1/CTNS-1) in dynamic CTNS-1 compartments in wild type, *sam-4(js415), arl-8(jpn1), sam-4(js415); arl-8(jpn1)*. Number of animals ≥ 12, Number of compartments ≥ 57, Statistical test : Kruskal Wallis ANOVA with Dunns test. F) Quantification of the area of dynamic CTNS-1 compartments in wild type, *sam-4(js415), arl- 8(jpn1), sam-4(js415); arl-8(jpn1)*. Number of animals ≥ 12, Number of compartments ≥ 57, Statistical test : Kruskal Wallis ANOVA with Dunns test. All data plotted as violin plots showing individual data points marked along with the median (solid line), 25th and 75th percentile marked (dashed lines). * p <0.05, ** p <0.01; *** p <0.001, ns - not significant. All relevant statistical comparisons are highlighted.

Additionally, *arl-8(lof)* mutants exhibited reduced CTNS-1 compartment area, similar to *sam- 4(0)* phenotypes (Fig 2A and 2C).

To test if SAM-4 acts through ARL-8 to segregate SVPs and lysosomal proteins and to regulate the size of LPCCs, we analyzed *sam-4(0); arl-8(gof)* double mutants. In *sam-4(0); arl-8(gof)* animals, we observed decreased Pearson’s and Mander’s colocalisation coefficients compared to *sam-4(0)*, suggesting improved segregation of SVPs and lysosomal proteins (S2C and S2D, Movie 11 and 12). The MFI ratio (SNG-1/CTNS-1) in moving CTNS-1 compartments was reduced in *sam-4(0); arl-8(gof)* compared to *sam-4(0)*, indicating improved segregation of SVPs and lysosomal proteins (Fig 2D, 2E; Movie 11 and 12). Interestingly, *arl-8(gof)* animals show an increased ratio of MFI (SNG-1/CTNS-1) but did not cause lysosome fragmentation indicating that segregation and fragmentation are separable processes (Fig 2D, 2E; Movie 11). Additionally, *arl-8(gof)* reversed *sam-4(0)* fragmentation defects, as shown by increased CTNS-1 compartment area (Fig 2D and 2F).

Together, these results suggest that BORC acts through ARL-8 to regulate SVP and lysosomal protein segregation and LPCC size. Furthermore, our findings indicate that fragmentation and segregation can be mechanistically distinct processes.

### UNC-104 regulates SVP-lysosomal protein segregation and UNC-116 regulates SVP- lysosomal protein segregation and the size of CTNS-1 compartments

ARL-8 can recruit UNC-104/KIF1A, KIF5B and Dynein microtubule motors [9,19,21–23,25,26]. ARL-8 recruits Kinesin-1 and Kinesin-3 through SKIP on lysosomes [21,26]. ARL-8 also regulates Dynein-driven retrograde lysosomal transport through RUFY3, RUFY4 and DENND6A [22,23,27]. ARL-8 is thought to activate the Kinesin-3 motor, UNC-104, thereby recruiting it to synaptic vesicles [25]. Since we observe defects in the segregation of SVPs and lysosomal proteins in BORC and ARL-8 mutants, we asked if these segregation defects arise from the inability to recruit any of the ARL-8 dependent motors.

We assessed the overlap of SNG-1::mNeonGreen and CTNS-1::mCherry in PLM cell bodies of the loss of function mutants *unc-104(e1265), unc-116(rh24sb79)* and *dhc-1(js319)* that encode the worm orthologues of KIF1A, KIF5 and Dynein. We observed an increase in the overlap of SNG-1 and CTNS-1, indicated by an increase in the Pearson’s coefficient in two alleles each of *unc-104* and *unc-116* mutants (Fig 3A, 3B; S3A and S3B, Movie 15 and 16). We did not observe a significant change in the overlap of SNG-1 and CTNS-1 in *dhc-1* mutant animals (Fig 3A and 3B). These data suggest that UNC-104/KIF1A and UNC-116/KIF5 have roles in the segregation of SVPs and lysosomal proteins. Further, we observed a significant increase in the ratio of the MFI (SNG-1/CTNS-1) in *unc-104* mutants (Fig 3C, 3D; Movie 17). However, the area of these dynamic compartments in *unc-104* mutants was not altered (Fig 3C and 3E). These data suggest that the segregation of SVPs and lysosomal proteins depends on *unc-104*, whereas the regulation of CTNS-1 compartment size is independent of *unc-104*.

**Figure 3:**
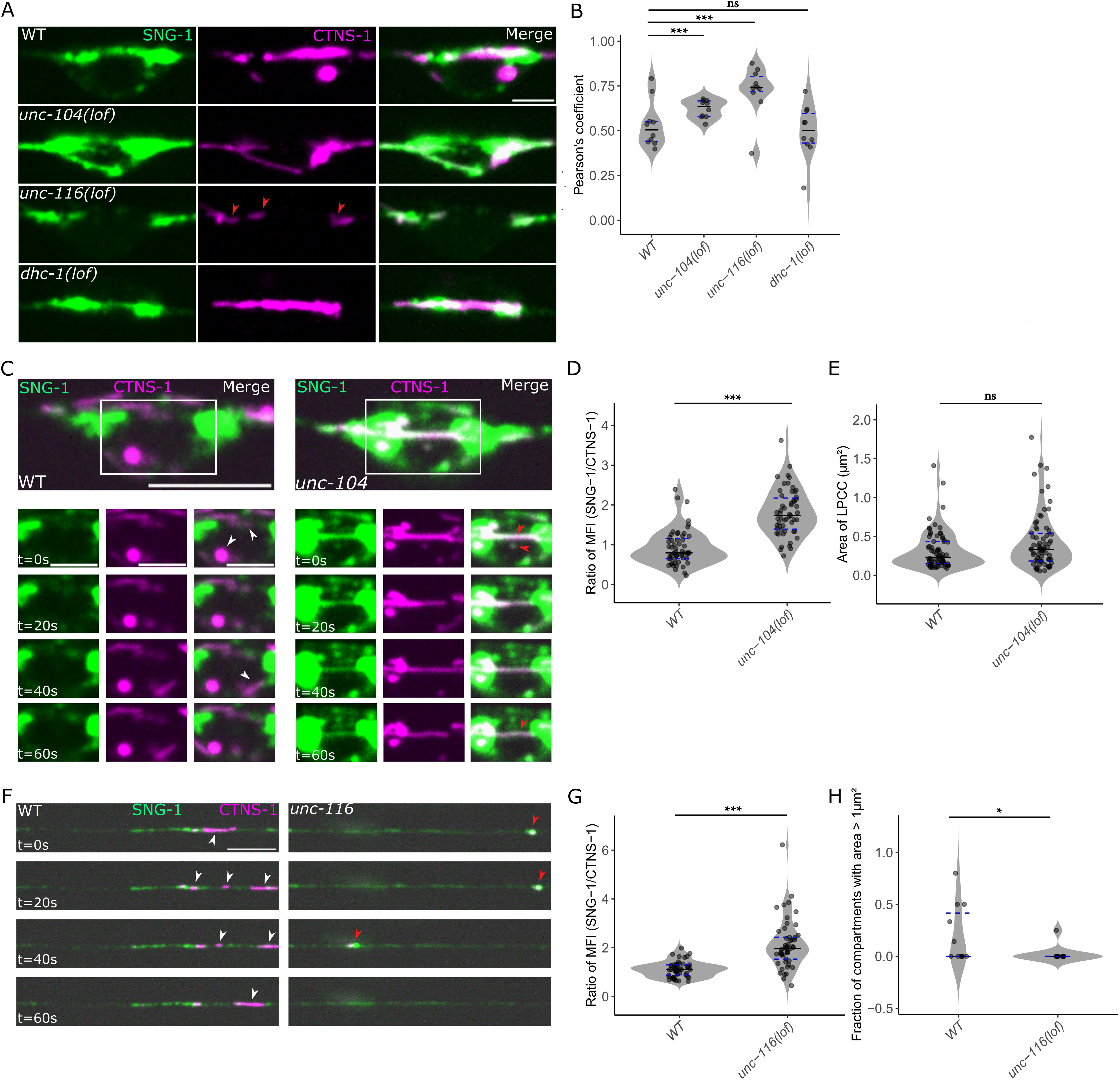
**UNC-104/KIF1A and UNC-116/KIF5 segregates SVPs and lysosomal proteins and UNC-116 additionally regulates the size of CTNS-1 compartments** A) Representative images of wild type, *unc-104(e1265), unc-116(rh24sb79)* and *dhc-1(js319)* showing the overlap of SNG-1::mNeonGreen (Green) and CTNS-1::mCherry (Magenta) in the PLM cell body. Red arrowheads indicate fragmented CTNS-1 compartments. Scale bar = 2.5 µm B) Quantification of the overlap of SNG-1 and CTNS-1 by Pearson’s Coefficient in wild type, *unc-104(e1265), dhc-1(js319)* and *unc-116(rh24sb79)*. Number of animals ≥ 10, Statistical test : Wilcoxon rank-sum test. C) Representative images of wild type and *unc-104(e1265)* showing the overlap of SNG- 1::mNeonGreen (Green) and CTNS-1::mCherry (Magenta) in dynamic compartments in the PLM cell body at time points t=0s to t=60s. Left set of panels corresponds to wild type and right set of panels corresponds to *unc-104(e1265)*. White arrowheads indicate compartments showing reduced SNG-1 with CTNS-1 while red arrowheads indicate compartments showing higher levels of SNG-1 in CTNS-1 compartments. Scale bar = 5 µm (cell body panel with ROI), 2.5 µm (time series panels). D) Quantification of the ratio of Mean Fluorescence Intensity (SNG-1/CTNS-1) in dynamic CTNS-1 compartments in wild type and *unc-104(e1265)*. Number of animals ≥ 13, Number of compartments ≥ 70, Statistical test : Wilcoxon rank-sum test. E) Quantification of the area of dynamic CTNS-1 compartments in wild type and *unc-104(e1265)*. Number of animals ≥ 13, Number of compartments ≥ 70, Statistical test : Wilcoxon rank-sum test. F) Representative images of wild type and *unc-116(rh24sb79)* showing the overlap of SNG- 1::mNeonGreen (Green) and CTNS-1::mCherry (Magenta) in dynamic compartments in the PLM minor neurite at time points t=0s to t=60s. Panels on the left correspond to wild type and panels on the right correspond to *unc-116(rh24sb79)*. White arrowheads indicate compartments showing reduced SNG-1 with CTNS-1 while red arrowheads indicate compartments showing higher levels of SNG-1 in CTNS-1 compartments. Scale bar = 5 µm. G) Quantification of the ratio of Mean Fluorescence Intensity (SNG-1/CTNS-1) of dynamic CTNS-1 compartments in the minor neurite of wild type and *unc-116(rh24sb79)*. Number of animals ≥ 11, Number of compartments ≥ 50, Statistical test : Wilcoxon rank-sum test. H) Quantification of the fraction of CTNS-1 compartments/LPCCs with area > 1 µm2 of dynamic CTNS-1 compartments in the minor neurite of wild type and *unc-116(rh24sb79)*. Number of animals ≥ 11, Number of compartments ≥ 50, Statistical test : Wilcoxon rank-sum test. All data plotted as violin plots showing individual data points marked along with the median (solid line), 25th and 75th percentile marked (dashed lines). * p <0.05, ** p <0.01; *** p <0.001, ns - not significant. All relevant statistical comparisons are highlighted.

*unc-116* mutants did not contain dynamic CTNS-1 compartments in the neuronal PLM cell body and hence we analyzed the ratio of MFI (SNG-1/CTNS-1) and area of dynamic CTNS-1 compartments in the minor neurite of PLM (S1A, Movie 18). We observed a significant increase in the ratio of MFI (SNG-1/CTNS-1) of CTNS-1 compartments in *unc-116(lof)* mutants (Fig 3F, 3G; Movie 19 and 20). We also observed a non-significant decrease in the size of CTNS-1 compartments in the minor neurite, however tubular CTNS-1 compartments were largely absent in *unc-116* mutants in the PLM minor neurite (S3C). Additionally, we observed a decrease in the fraction of animals that have LPCC compartments with area greater than 1μm² in *unc-116* mutants in the PLM minor neurite (Fig 3F and 3H).

Together, we have identified a novel role for the Kinesin-3 motor, UNC-104, and the Kinesin-1 motor, UNC-116, in segregating SVPs and lysosomal proteins and an additional role for UNC- 116 in regulating the area of CTNS-1 compartments.

To test if SAM-4 and ARL-8 act through the motors, we quantified the Pearson’s correlation coefficient for SNG-1::mNeonGreen and CTNS-1::mCherry in the triple mutants *sam-4(0); unc- 116(lof); arl-8(gof)* and *sam-4(0) unc-104(lof); arl-8(gof)*. We observed that the Pearson’s correlation coefficient *sam-4(0); unc-116(lof); arl-8(gof)* was significantly different from *sam- 4(0); arl-8(gof)* (Fig S5A and S5B). This indicates that the segregation of SVPs & lysosomal proteins by SAM-4 & ARL-8 depends on UNC-116/KIF5A. We also observed that the Pearson’s correlation coefficient for SNG-1 and CTNS-1 in PLM cell bodies of *sam-4(0) unc-104(lof); arl- 8(gof)* was closer to *sam-4(0)* (p=0.2) than *sam-4(0); arl-8(gof)* (p=0.08) (Fig S6A and S6B).

This suggests that the improved segregation of SNG-1 and CTNS-1 in *sam-4(0); arl-8(gof)* neuronal cell bodies is partially dependent on UNC-104.

Together, our data indicates that BORC and ARL-8 act through motor proteins in the genetic pathway for SVP-lysosomal protein segregation and regulating size in the neuronal cell body. Additionally, the ability of constitutively active ARL-8 to enable segregation of SVPs and lysosomal proteins in the absence of SAM-4 depends on its ability to recruit both UNC-116 and UNC-104. Since *unc-104* mutants only exhibit a subset of phenotypes shown by *unc-116* mutants, UNC-116 may act before UNC-104 in the sorting pathway.

### BORC acts through ARL-8 and UNC-104 to regulate the distribution of SV-lysosomes in the axon

Lysosomal markers and specifically CTNS-1 compartments where the majority that exit the cell body carry SVPs (SV-lysosomes) are present just in the proximal axons of *C. elegans* [8,17]. The extent of lysosomal protein containing compartments present along the axon is dependent on UNC-104/KIF1A [8,11]. We thus investigated the role of BORC, ARL-8, and the Kinesin motors in regulating the distribution of CTNS-1-containing compartments in axons.

First, we assessed the distance that CTNS-1::mCherry travels into axons of PLM neuron in *sam- 4(0)* and *arl-8(lof)* mutants. We observed a significant decrease in the distance CTNS-1 compartments was present along the PLM major neuronal process/axon in both *sam-4(0)* and *arl-8(lof)* (Fig 4A, 4B, S6A, S6B and S6C). Mutants in *snpn-1*, another component of the BORC complex, also showed reduced distance that CTNS-1 compartments was present along the axon while mutants in *dsbn-1*, a component of the BLOC-1 complex, did not show any change in the distribution of CTNS-1 compartments in the neuronal process (Fig 4A and 4B). Next, we asked whether the gain-of-function (gof) allele, *arl-8(jpn1)*, a GTP-locked constitutively active ARL-8, could reverse the restricted CTNS-1 compartment distribution observed in *sam-4* mutants. We observed an increased exit of CTNS-1 compartments into the axon in *sam-4(0); arl-8(gof)* double mutants indicating that SAM-4 acts through ARL-8 in regulating the distribution of CTNS-1 compartments (Fig 4C, 4D).

**Figure 4:**
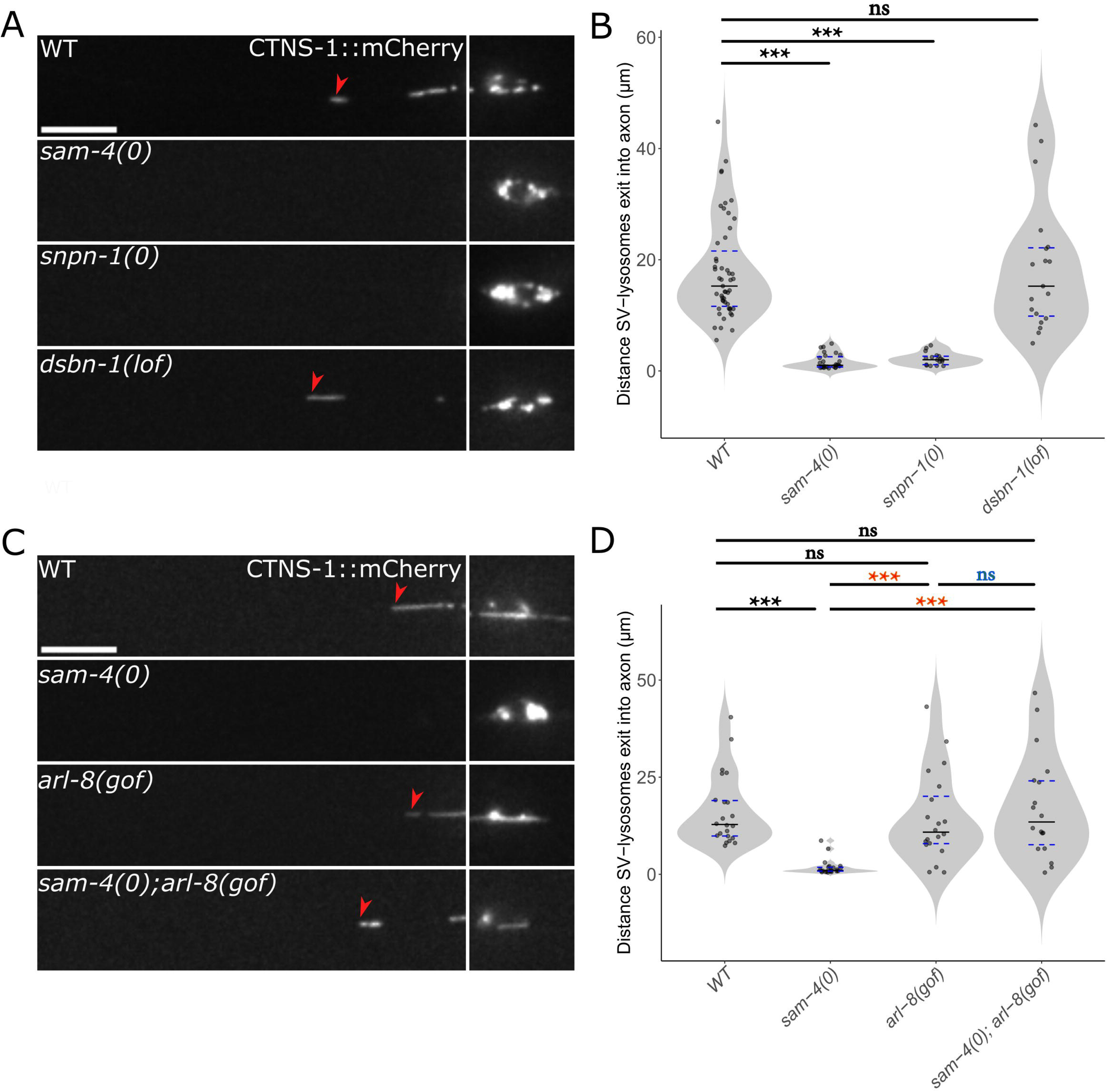
BORC through ARL-8 and UNC-104/KIF1A regulates the distribution of SV- lysosomes in *C. elegans* TRNs A) Representative images of wild type, *sam-4(js415), snpn-1(tm1892)* and *dsbn-1(tm6580)* showing the distribution of CTNS-1::mCherry. Red arrowhead indicates the distance CTNS-1 exits into the neuronal process. Scale bar = 10 µm. B) Quantification of the distance CTNS-1 exits into the neuronal process in wild type, *sam- 4(js415), snpn-1(tm1892)* and *dsbn-1(tm6580).* Number of animals ≥ 17, Statistical test : Wilcoxon rank-sum test C) Representative images of wild type, *sam-4(js415), arl-8(jpn1)* and *sam-4(js415); arl-8(jpn1)* showing the distribution of CTNS-1::mCherry. Red arrowhead indicates the distance CTNS-1 exits into the neuronal process. Scale bar = 10 µm D) Quantification of the distance CTNS-1 exits into the neuronal process in wild type, *sam- 4(js415), arl-8(jpn1)* and *sam-4(js415); arl-8(jpn1).* Number of animals ≥ 18, Statistical test : Kruskal Wallis ANOVA with Dunns test. All data plotted as violin plots showing individual data points marked along with the median (solid line), 25th and 75th percentile marked (dashed lines). * p <0.05, ** p <0.01; *** p <0.001, ns - not significant. All relevant statistical comparisons are highlighted.

We further investigated the role of motor proteins in regulating CTNS-1 compartment distribution along the neuronal process. In *unc-104(lof)* mutants, we observed a reduction in the distance CTNS-1 containing compartments traveled into the axon (S6D, S6E, S7A and S7B). In *unc-116(lof)* mutants, CTNS-1 compartments are mislocalised to minor neurites and the distal tips of major processes (S6D and S6E). These data suggest that the exit of CTNS-1 compartments into the axon depends on UNC-104. In *unc-116* mutants, mislocalisation of CTNS-1 compartments to minor neurite likely results from altered microtubule polarity, while mislocalisation to the distal tip of the major process is likely due to the role of UNC-116 in dynein transport [28–30]. Consistent with this hypothesis we observe that CTNS-1 compartments are present up to a greater distance in the axon in *dhc-1* mutants compared to wild type (S6F and S6G).

Finally, we examined the role of UNC-104 in the exit of CTNS-1 compartments into the axon by assessing the triple mutant, *sam-4(0) unc-104(lof); arl-8(gof)* mutants. The reduced CTNS-1 compartment exit into axons observed in these mutants indicates that the exit of CTNS-1 compartments in *sam-4(0) arl-8(gof)* mutants depends on UNC-104 (S7A and S7B).

ARL-8 is known to recruit motor proteins and our data suggests that BORC through ARL-8 and UNC-104 regulates the entry of CTNS-1 compartments that are likely SV-lysosomes in the axon potentially through the ability to recruit UNC-104. The distribution of CTNS-1 compartments along neuronal processes depends on both UNC-116 and dynein motors as well.

### LRRK2/LRK1 acts through BORC to regulate synaptic vesicle protein and lysosomal protein segregation and size of CTNS-1 compartments

LRK-1/LRRK2 plays a key role in lysosomal and synaptic vesicle protein trafficking [7,8,31–33]. *lrk-1* mutants show an increased presence of SV-lysosomal compartments and increased size of SVP carriers in the axon [7,8]. We therefore asked (i) Do *lrk-1* mutants show defective segregation of SVPs and lysosomal proteins in the cell body? (ii) If yes, does this segregation depend on BORC-ARL-8? (iii) Does LRK-1 affect BORC? We assessed the overlap of SNG-1::mNeonGreen and CTNS-1::mCherry in the PLM cell bodies of the loss of function mutant *lrk-1(km17)* that carries a deletion in its kinase domain [31]. *lrk- 1(lof)* animals show a significant increase in the overlap of SNG-1 and CTNS-1 in the PLM cell body as indicated by the Pearson’s and Mander’s correlation coefficients (Fig 5A; S8A, S8B and Movie 21). We also observed an increased ratio of MFI (SNG-1/CTNS-1) in *lrk-1(lof)* mutants indicating an increased presence of SVPs in moving CTNS-1 compartments (Fig 5A, 5B; Movie 22). *lrk-1(lof)* mutants also showed reduced area of CTNS-1 compartments similar to *sam-4(0)* (Fig 5A and 5C). These data suggest that *lrk-1(lof)* and *sam-4(0)* both regulate the segregation of SVPs and lysosomal proteins and size of LPCCs in the neuronal cell body.

**Figure 5:**
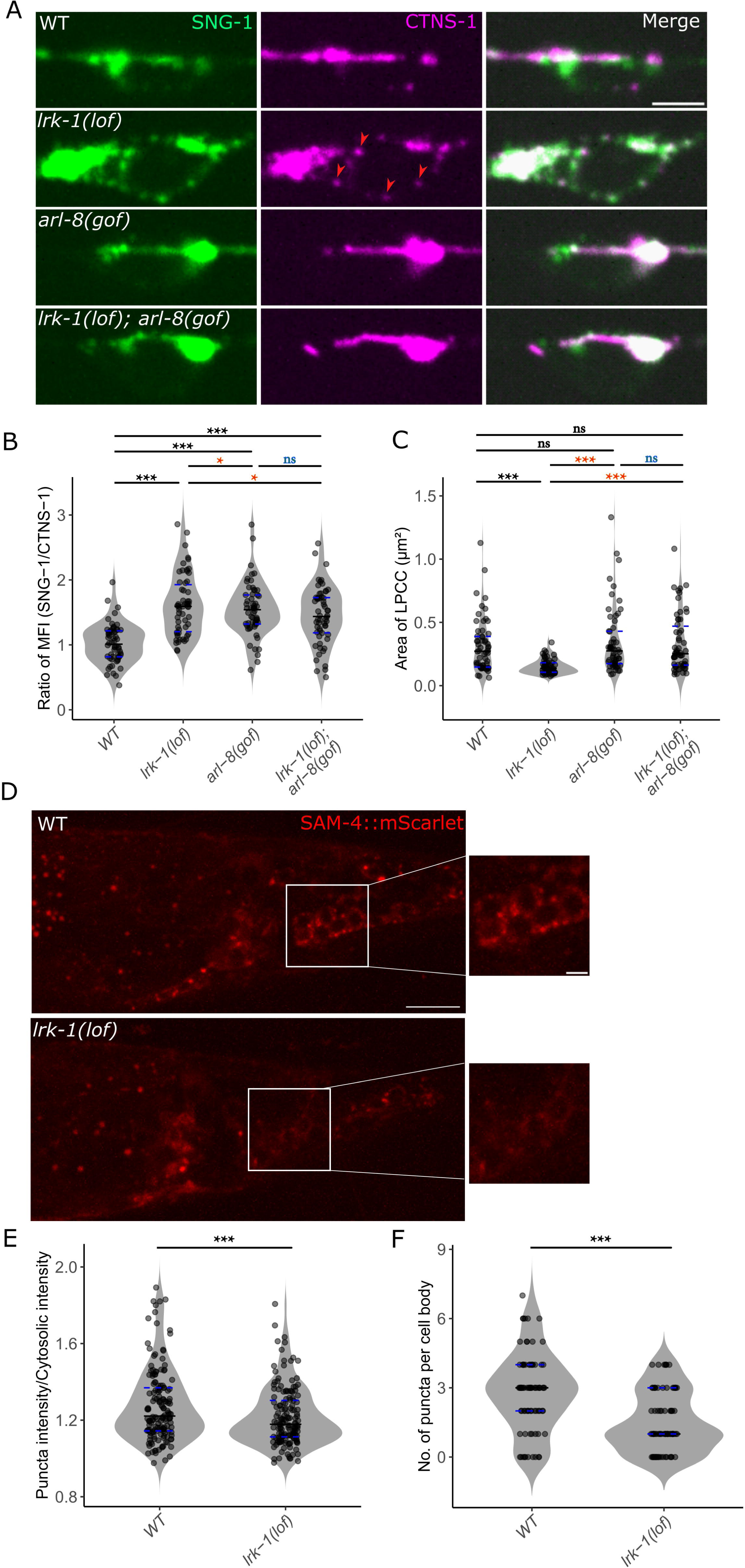
**LRK-1 through BORC segregates SVPs and lysosomal proteins and regulates the size of CTNS-1 compartments** A) Representative images of wild type, *lrk-1(km17), arl-8(jpn1)* and *lrk-1(km17); arl-8(jpn1)* showing the overlap of SNG-1::mNeonGreen (Green) and CTNS-1::mCherry (Magenta) in the PLM cell body. Red arrowheads indicate fragmented CTNS-1 compartments. Scale bar = 2.5 µm B) Quantification of the ratio of Mean Fluorescence Intensity (SNG-1/CTNS-1) in dynamic CTNS-1 compartments in wild type, *lrk-1(km17), arl-8(jpn1)* and *lrk-1(km17); arl-8(jpn1)*. Number of animals ≥ 11, Number of compartments ≥ 58, Statistical test : Kruskal Wallis ANOVA with Dunns test. C) Quantification of the area of dynamic CTNS-1 compartments in wild type, *lrk-1(km17), arl- 8(jpn1)* and *lrk-1(km17); arl-8(jpn1)*. Number of animals ≥ 11, Number of compartments ≥ 58, Statistical test : Kruskal Wallis ANOVA with Dunns test. D) Representative images of wild type and *lrk-1(km17)* showing the localisation of SAM- 4::mScarlet (red) in the cell bodies at the tail of *C. elegans*. Scale bar =10 µm (zoomed out panel), 2.5 µm (zoomed in panel) E) Quantification of the intensity (Puncta intensity/Cytosolic intensity) of SAM-4 puncta from the tail cell bodies in wild type and *lrk-1(km17).* Number of animals ≥ 10, Number of cell bodies ≥ 81, Number of puncta ≥ 165, Statistical test : Wilcoxon rank-sum test. F) Quantification of the number of SAM-4 puncta per cell body from the tail cell bodies in wild type and *lrk-1(km17).* Number of animals ≥ 10, Number of cell bodies ≥ 81, Number of puncta ≥ 165, Statistical test : Wilcoxon rank-sum test. All data plotted as violin plots showing individual data points marked along with the median (solid line), 25th and 75th percentile marked (dashed lines). * p <0.05, ** p <0.01; *** p <0.001, ns - not significant. All relevant statistical comparisons are highlighted.

We further asked if constitutively active *arl-8(jpn1)*, that can recruit multiple motors that control segregation & compartment size, reverses the segregation and fragmentation defects in *lrk-1(lof)*. We observed a significant reduction in the ratio of MFI (SNG-1/CTNS-1) of moving compartments in *lrk-1(lof); arl-8(gof)* compared to *lrk-1(lof)* indicating that constitutively active ARL-8 that can recruit multiple motors can improve segregation of SVPs and lysosomal proteins and reverse fragmentation defects in *lrk-1(lof)* (Fig 5A, 5B; Movie 23). However, the overlap of SNG-1 and CTNS-1 as assessed by Pearson’s and Mander’s correlation coefficients in *lrk-1(lof); arl-8(gof)* shows only a small but non-significant reduction compared to *lrk-1(lof)* (S8A, S8B and Movie 24). We also observed that *arl-8(gof)* reversed the fragmentation defects in *lrk-1(lof)* as observed by an increase in the area of compartments in *lrk-1(lof); arl-8(gof)* compared to *lrk- 1(lof)* (Fig 5A and 5C). These data suggest that LRK-1 can regulate SVP and lysosomal protein segregation and LPCC size. Additionally, LRK-1 might act through ARL-8 in mediating these functions.

We assessed the localisation of SAM-4 in the loss of function mutants of *lrk-1(km17)*. SAM-4 is known to be associated with membranes and in wild type cell bodies SAM-4 shows a punctate localisation [16,17]. We observed a significant reduction in the intensity of puncta and the number of puncta of SAM-4::mScarlet (CRISPR knock-in) in *lrk-1(lof)* mutants (Fig 5D, 5E and 5F) [24]. Additionally, we also observe that the intensity and the localisation of SAM-4 puncta is altered in *snpn-1(0)* mutants indicating that loss of one protein in the BORC complex may alter the localisation of other proteins in the complex (S9A).

These findings suggest that LRK-1 can regulate the segregation of SVPs and lysosomal proteins and the size of LPCCs in the cell body by acting through ARL-8. Additionally, LRK-1 governs these functions by mediating the correct localisation of BORC.

### The clathrin adaptor complex AP-3, regulated by BORC, segregates synaptic vesicle proteins and lysosomal proteins

The AP-3 complex of *C. elegans* consists of the gene products encoded by the *apm-3* (µ3), *apb-3* (β3), *apd-3* (δ) and *aps-3* (σ3) genes [34]. This adaptor complex plays key roles in lysosome biogenesis and also regulates the trafficking of SVPs [8,12,35]. *apb-3* mutants show an increased presence of SV-lysosomal compartments in the neuronal process and LRK-1 is thought to act via APB-3 to facilitate SV membrane composition and SV-lysosomal presence in axons [7,8]. Since phenotypes exhibited by *apb-3* mutants overlap with *lrk-1* mutants, we asked i) Do *apb-3* mutants show defective segregation of SVPs and lysosomal proteins in the cell body? ii) Does APB-3 regulate BORC localisation and/or vice-versa? We examined the overlap of SNG-1::mNeonGreen and CTNS-1::mCherry in PLM cell bodies of *apb-3(ok429),* a null deletion mutant [36]. Mutants in *apb-3(0)* show a significant increase in the overlap of SNG-1::mNeonGreen and CTNS-1::mCherry in the PLM neuronal cell body as indicated by the Pearson’s and Mander’s correlation coefficients and an increased MFI ratio (SNG-1/CTNS-1) (Fig 6A, 6B; S8C, S8D, Movie 25 and 26). Unlike *sam-4(0)* and *lrk-1(lof)* animals, *apb-3(0)* mutants do not show a reduced CTNS-1 compartment size (Fig 6A and 6C).

**Figure 6:**
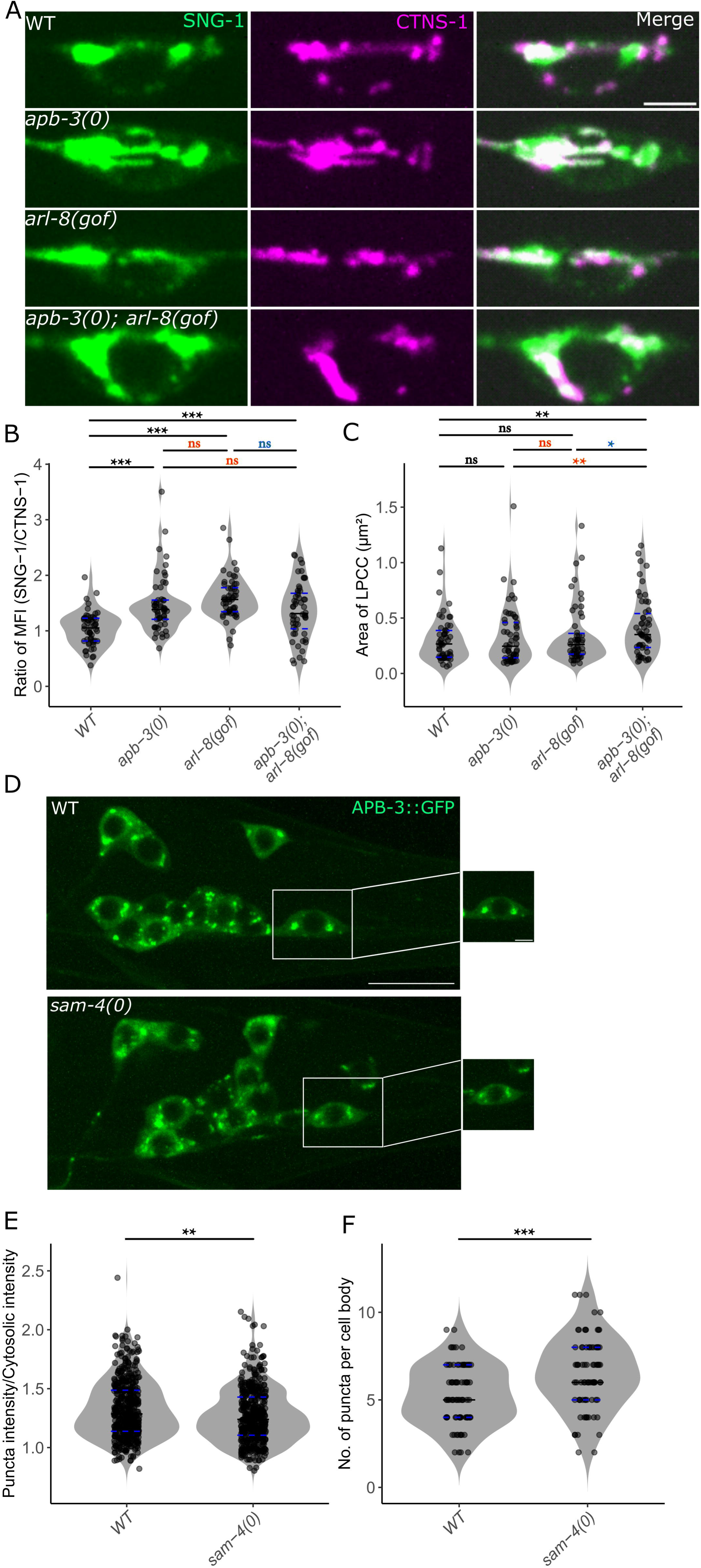
**BORC acts through APB-3 in segregating SVPs and lysosomal proteins** B) Representative images of wild type, *apb-3(ok429), arl-8(jpn1)* and *apb-3(ok429); arl-8(jpn1)* showing the overlap of SNG-1::mNeonGreen (Green) and CTNS-1::mCherry (Magenta) in the PLM cell body. Scale bar = 2.5 µm. C) Quantification of the ratio of Mean Fluorescence Intensity (SNG-1/CTNS-1) in dynamic CTNS-1 compartments in wild type, *apb-3(ok429), arl-8(jpn1)* and *apb-3(ok429); arl-8(jpn1)*. Number of animals ≥ 12, Number of compartments ≥ 54, Statistical test : Kruskal Wallis ANOVA with Dunns test. D) Quantification of the area of dynamic CTNS-1 compartments in wild type, *apb-3(ok429), arl- 8(jpn1)* and *apb-3(ok429); arl-8(jpn1)*. Number of animals ≥ 12, Number of compartments ≥ 54, Statistical test : Kruskal Wallis ANOVA with Dunns test. E) Representative images of wild type and *sam-4(js415)* showing the localisation of APB-3::GFP (green) in the cell bodies at the tail of *C. elegans*. Scale bar =10 µm, 2.5 µm (zoomed in panel). F) Quantification of the intensity (Puncta intensity/Cytosolic intensity) of APB-3::GFP puncta from the tail cell bodies in wild type and *sam-4(js415).* Number of animals ≥ 10, Number of cell bodies ≥ 99, Number of compartments ≥ 514, Statistical test : Wilcoxon rank-sum test. G) Quantification of the number of APB-3::GFP puncta per cell body from the tail cell bodies in wild type and *sam-4(js415).* Number of animals ≥ 10, Number of cell bodies ≥ 99, Number of puncta ≥ 514, Statistical test : Wilcoxon rank-sum test. All data plotted as violin plots showing individual data points marked along with the median (solid line), 25th and 75th percentile marked (dashed lines). * p <0.05, ** p <0.01; *** p <0.001, ns - not significant. All relevant statistical comparisons are highlighted.

These results suggest that *apb-3(0)* shares phenotypes with *sam-4(0)* and *lrk-1(lof)* in regulating SVP and lysosomal protein segregation, but not in LPCC fragmentation.

We further tested whether the *arl-8(jpn1)* can improve the segregation of SVPs and lysosomal proteins in *apb-3(0)*. The overlap of SNG-1 and CTNS-1 quantified by Mander’s coefficient and ratio of MFI (SNG-1/CTNS-1) showed a non-significant reduction in *apb-3(0); arl-8(gof)* compared to *apb-3(0)*, Pearson’s coefficient showed a significant reduction (Fig 4B; S8C, S8D, Movie 27 and 28). Since the ratio of MFI (SNG-1/CTNS-1) and Mander’s do not show a significant reduction we think that the small reduction in Pearson’s correlation coefficient is likely due to partial reduction in the levels of SVPs in static CTNS-1 containing compartments. Together, these findings suggest SVP and lysosomal protein segregation by APB-3 might not completely depend on ARL-8.

Further, we assessed the localisation of APB-3 and SAM-4 in *sam-4(0)* and *apb-3(0)* mutants respectively. We did not observe a change in the intensity of puncta and the number of puncta of SAM-4 in *apb-3(0)* mutants (S9A, S9B and S9C). Furthermore, we tested if SAM-4 regulates APB-3 localisation. Approximately, 66% of APB-3 puncta juxtapose with CTNS-1 (S10A, S10B, Movie 29). We observed a significant decrease in the intensity of APB-3 puncta and an increase in the number of APB-3 puncta per cell body in *sam-4(0)* mutants (Fig 6D, 6E and 6F). These findings suggest that SAM-4 may regulate APB-3 localisation.

Together, our data suggest that APB-3 can regulate the segregation of SVPs and lysosomal proteins in the cell body. Since APB-3 continues to show punctate localisation in *sam-4(0)* mutants, we think that SAM-4 does not affect recruitment to membranes but regulates the levels of APB-3 on sorting compartments.

## Discussion

Sorting synaptic vesicle proteins away from lysosomal proteins is one of the steps in the biogenesis of pre-SVs and lysosomes [8,12,14]. We have uncovered a novel role for BORC acting through ARL-8 and motor proteins in segregating SVPs and lysosomal proteins and regulating the size of LPCCs in *C. elegans* neuronal cell body. Segregation of SVPs and lysosomal proteins and size regulation by BORC through ARL-8 is linked with the action of motors UNC-116 and UNC-104. Recruitment of motors could mediate protein segregation and tubulation from sorting compartments. Additionally, we show that LRK-1 acts through BORC and through APB-3 to mediate the segregation of SVPs and lysosomal proteins in the cell body.

Although SV-lysosomal compartments have been seen in axons, it is not clear if these compartments are biosynthetic compartments. A recent study using the RUSH system showed that biosynthetic LAMP2-containing compartments are co-transported in hippocampal neurons with synaptic vesicle proteins like synaptotagmin, consistent with the idea that SV-lysosomes in the axons can be biosynthetic compartments prior to segregation of synaptic vesicle and lysosomal proteins [14]. Additionally, our data indicate that the sorting events at the cell body mediated by LRK-1, BORC, ARL-8 and APB-3 could account for the non-overlap of SVP and lysosomal proteins in mouse hippocampal neurons and also fewer SV-lysosomes in axons in *C. elegans*.

LRRK2/LRK-1 is a master regulator of membrane trafficking and it phosphorylates several proteins involved in membrane trafficking including RAB GTPases [37–39]. Casein Kinase Ια- like isoform has been shown to regulate the membrane localisation of AP-3 through phosphorylation [40]. Given that LRK-1 influences BORC localisation, it is possible that BORC may be phosphorylated by LRK-1. LRK-1-mediated regulation of BORC localisation could play a critical role in controlling the segregation and size of lysosomal protein-containing compartments (LPCCs) (Fig. 5D–F). Identifying downstream targets of LRK-1 is of great interest, as mutations in LRK-1 are implicated in Parkinson’s disease. Our study identifies BORC as a downstream effector of LRK-1, which could have significant implications. The LRK-1-BORC pathway could also act in parallel with LRK-1-APB-3 pathway to mediate the segregation of SVPs and lysosomes or LRK-1 could act through BORC and further through or additionally through APB-3 to mediate the segregation of SVPs and lysosomal proteins in the cell body. Additionally, our study highlights the need to assess BORC’s role in the neuronal cell body of higher model organisms.

Our study also identifies a role for BORC in regulating the size of lysosomal protein-containing compartments (LPCCs) (Fig 1F). BLOC-1 which shares subunits with BORC is known to tubulate endosomal membranes [41,42]. This regulation might result from defective recruitment of UNC-116 to SV-lysosomal compartments or other membrane-modifying proteins (Fig 4F). Motor proteins are known to tubulate compartments [43–45]. Tubulation can occur as a consequence of force generation on membranes resulting from microtubule binding and walking [43,44,46]. Additionally, our observations are consistent with a recent study that shows that BORC through ARL-8 and UNC-116 tubulates phagocytic cargo and mediates their clearance [24].

We propose that tubulation may facilitate sorting by clustering specific proteins or lipids through the cargo-binding domains of motors, leading to segregation based on affinity. While Dynein is implicated in cargo sorting, the role of Kinesins remains unexplored [47,48]. We hypothesize that UNC-104 and UNC-116 mediate the segregation of SVPs and lysosomal proteins by binding to SV-lysosomal compartments and sorting proteins based on the affinity of their cargo-binding domains. This model supports at least three possibilities i) pre-SVs are formed by active sorting of proteins and lipids as a consequence of motor binding on SV-lysosomal intermediates ii) pre-SVs are remnants of SV-lysosomal compartments after lysosomal proteins are sorted away iii) active sorting by two sets of motors/tubulation proteins acts mutually exclusively on SV-lysosomal compartments to sort SVPs and lysosomal proteins away from each other. AP-3, a key adaptor protein involved in lysosome, lysosome-related organelles (LROs), and synaptic vesicle biogenesis, may also contribute by sorting transmembrane proteins [7,49–51]. Since pre-SVs contain UNC-104, we propose that UNC-104 and AP-3 act mutually exclusively on SV-lysosomal intermediates, with AP-3 sorting lysosomal proteins and UNC-104 sorting SVPs, potentially in concert with other adaptors. This model explains the observed sorting phenotypes of UNC-104/UNC-116 and AP-3.

In conclusion, our data suggest that BORC, likely acting through motor proteins UNC-116 and UNC-104, regulates the segregation and size of lysosomal compartments. This regulation likely occurs at SV-lysosomal intermediates, where AP-3 and motor proteins regulated by BORC mediate segregation and size control. BORC is regulated by the Parkinson’s-associated gene LRK-1. Together, these processes and genetic pathways play a crucial role in the biogenesis of pre-SVs.

## Acknowledgments

We thank Gholamreza Fazeli and Ann Wehmann for providing us with SAM-4::mScarlet (CRISPR knock-in) strain. We thank Shraddha Athawale for building and imaging *dhc-1(js319); tbIs381*. Research in Sandhya Koushika’s lab is supported by grants from the Department of Atomic Energy, Government of India (DAE; OM no. 1303/2/2019/R&D-II/DAE/2079; Project identification number RTI4003 dated 11.02.2020). Research in Michael L Nonet’s Lab is supported by a National Institutes of Health research grant (R01 GM14168802).

## Materials and Methods

### Strain maintenance

*C. elegans* were grown on nematode growth media (NGM) spotted with *Escherichia coli* OP50 culture at 20°C. Preparation of NGM plates and culturing of OP50 was done following standard protocols from Wormbook [52]. All strains used in this study are listed in Table S1.

### Creation of transgenic lines

Insertions were created using RMCE, rRMCE or RMHE. Injections were performed as described in (Nonet, 2023). Most animals were only injected in a single gonad. DNAs were injected at ∼ 50 µg/ml in 10 mM Tris pH 8.0, 0.1 mM EDTA. RMCE was performed using a *sqt-1* Rol screening strategy [53]. For rRMCE using drug selection, injected P0 animals were pooled 2-3 per plate in most cases [54]. Hygromycin B (GoldBio, St. Louis, MO; 100 µl of 20 mg/ml) was added directly to worm plates 3 days after injection. Eight days after injection L4 animals homozygous for insertions were isolated from the injection plates. RMHE was performed by crossing *jsSi1784* phiC31 males into the *myo-2 nlsGFP marked attP* reporter line [55]. Cross progeny males were crossed to the unmarked *attB* driver line at 25° C to obtain reporter/driver *trans* heterozygotes also expressing the recombinase. The recombined driver::reporter transgene was identified among cross progeny of *trans* heterozygotes animals as animals expressing the reporter, but lacking the *myo-2p nlsGFP* marker *cis* linked to the reporter. The integration plasmid and parental strain used to construct each novel transgenic insertion and the strains containing these alleles are listed in Table S2.

### Imaging and Image Analysis Preparation of slides

L4 animals were anesthetized with 5 mM tetramisole or 10mM sodium azide and laid on a 5% agar pads. Imaging was done using two microscopes: i) an Olympus IX83 with a Yokogawa CSU-W1 and having a Prime BSI sCMOS camera and ii) an IX73 scope illuminated with a 120 W X-cite mercury-arc lamp having a Photometrics Evolve 512 EMCCD camera.

### Colocalisation analysis

Colocalisation analysis was performed on PLM cell body Z-stacks that cover the whole planes of the PLM cell body. We image 1 PLM cell body/animal. Confocal Z stacks were acquired using an Olympus IX83 fitted with a Yokogawa CSU-W1 at objective using a 100×/1.4 NA DIC oil objective and imaged with a Prime BSI sCMOS camera using 473 and 561 solid state lasers at a laser power of 3% and exposure of 250 ms for both channels. SoRA module in the cellSens software was chosen with Z slices taken at an interval of 0.28 µm and an effective pixel size of 0.065 um. The image files were opened and analysed using ImageJ. Colocalisation analysis was performed using the BIOP-JACOP plugin in ImageJ. ROI is drawn using the polygon selection tool around the cell body boundary. Pearson’s and Mander’s coefficient within the ROI is calculated using the BIOP-JACOP plugin. Mander’s coefficient is calculated using threshold images with the following thresholds used for each marker (SNG-1::mNeongreen - 300, CTNS- 1::mCherry - 200 , SNB-1::eGFP - 300, LAAT-1::mScarlet - 600). Pearson’s coefficient quantifies the correlation between the red and green channels and Mander’s coefficient (M2) quantifies the fraction of area in the red channel that overlaps with the green channel. For *unc- 116* mutants, ∼10% of animals do not have CTNS-1 signal in the cell body, such animals are not imaged for colocalisation analysis.

### Measuring Mean fluorescence intensity (MFI) and size of endolysosomal compartments

MFI and size measurements were done on dual-colour movies taken from the medial plane of the PLM cell body. We image 1 PLM cell body/animal. Dual colour movies were acquired using an Olympus IX83 fitted with a Yokogawa CSU-W1 at objective using a 100×/1.4 NA DIC oil objective and imaged with a Prime BSI sCMOS camera sequentially using 473 and 561 solid state lasers at a laser power of 5% and exposure of 200 ms for both channels. SoRA module in the cellSens software was chosen and the movie was taken at a frame rate of 2.3 frames/sec for 200 frames with an effective pixel size of 0.065 µm. The image files were opened and analysed using ImageJ. Dynamic lysosomal compartments were chosen for the analysis with a minimum distance cutoff of 3 pixels. These dynamic compartments were chosen from regions without stationary synaptic vesicle proteins, especially regions close to the Golgi. ROIs were drawn using the polygon selection tool in ImageJ and the average intensity within this ROI was measured using ImageJ. MFI (green)/MFI (red) was calculated by dividing the average intensities from both channels. Ratio of MFI (SNG-1/CTNS-1) quantifies the levels of SNG-1 in CTNS-1 compartments relative to CTNS-1 levels. Area was calculated using the same ROIs that were used for calculating the MFI values.

### Quantifying the number of puncta, area and intensity of APB-3::GFP and SAM- 4::mScarlet

Localisation of SAM-4 and APB-3 were quantified on Z-stacks that covered the whole tail. For APB::GFP, Confocal Z stacks were acquired using an Olympus IX83 fitted with a Yokogawa CSU-W1 at objective using a 100×/1.4 NA DIC oil objective and imaged with a Prime BSI sCMOS camera using 473 solid state laser at a laser power of 5% and an exposure of 400ms.

SoRA module in the CellSens software was chosen with Z slices taken at an interval of 0.28 µm and an effective pixel size of 0.065 µm. The image files were opened and analyzed using ImageJ. For SAM-4::mScarlet, Confocal Z stacks were acquired using an Olympus IX83 fitted with a Yokogawa CSU-W1 at objective using a 60×/1.35 NA DIC oil objective and imaged with a Prime BSI sCMOS camera using 561 solid-state laser at a laser power of 20% and an exposure of 500 ms. SoRA module in the cellSens software was chosen with Z slices taken at an interval of 0.33 µm and an effective pixel size of 0.108 µm. The image files were opened and analysed using ImageJ. ROIs were drawn around the punctas of APB-3 and SAM-4 in the tail cell bodies. For both APB-3::GFP and SAM-4::mScarlet, the number of puncta was calculated from all the planes covering each cell body. Using the ROI marked, the size and intensity of each puncta were measured. A cytosolic region close to the puncta was considered for background intensity. Puncta intensity was quantitated by dividing the intensity of each puncta by the cytosolic intensity. The same ROIs were also used to measure the area of punctas. All puncta values are plotted.

### Measuring distance SV-lysosomal compartments travel in the neuronal process

Distance endolysosomal compartments travel into the neuronal process was quantified from static images taken from an epifluorescence IX73 scope illuminated with 100% output from a 120 W X-cite mercury-arc lamp and imaged using a Photometrics Evolve 512 EMCCD camera and a 60×/1.35 NA DIC oil objective with a pixel size of 265 nm per pixel. An ROI is drawn from the edge of the cell body along the process to the end of the last endolysosomal compartment in the neuronal process. The length of this ROI is measured and plotted.

### Quantifying the fraction of APB-3 juxtaposing with CTNS-1

The juxtaposition of APB-3 with CTNS-1 was quantified using PLM cell body Z-stacks that cover the whole planes of the PLM cell body. We image 1 PLM cell body/animal. Confocal Z stacks were acquired using an Olympus IX83 fitted with a Yokogawa CSU-W1 at objective using a 100×/1.4 NA DIC oil objective and imaged with a Prime BSI sCMOS camera using 473 and 561 solid state lasers at a laser power of 10% and 3% respectively at an exposure of 500 ms for both channels. SoRA module in the cellSens software was chosen with Z slices taken at an interval of 0.28 µm and an effective pixel size of 0.065 µm. The image files were opened and analysed using ImageJ. The images were thresholded (APB-3::GFP - 300, CTNS-1::mCherry - 300) using the ImageJ plugin BIOP-JACOP. The thresholded images were used to visualize the overlap of APB-3 and CTNS-1. One puncta of APB-3 is considered to juxtapose with CTNS-1 if we observe at least 3 pixel overlap between APB-3 and CTNS-1. Each data point represents the fraction of APB-3 puncta juxtaposing with CTNS-1 per cell body.

### Statistical analysis

The Shapiro–Wilk test was used to test the normality of various distributions. For normally distributed data Unpaired two-tailed Student’s *t*-test was used for single comparisons and One-way ANOVA with Bonferroni test was used for multiple comparisons. For data that rejected normality, Wilcoxon rank-sum test was used for single comparisons and Kruskal Wallis ANOVA with Dunn’s Test was used for multiple comparisons. All data plotted as violin plots showing individual data points marked along with the median (solid line), 25th and 75th percentile marked (dashed lines). * p <0.05, ** p <0.01; *** p <0.001, ns - not significant. Table S3 contains all values used to generate all the graphs in the manuscript. Table S4 contains all the statistical comparisons in each figure.

