## Supplementary Figures for "BORC segregates synaptic vesicle and lysosomal proteins through motors UNC-104/KIF1A and UNC-116/KIF5"

Supplementary Figure 1

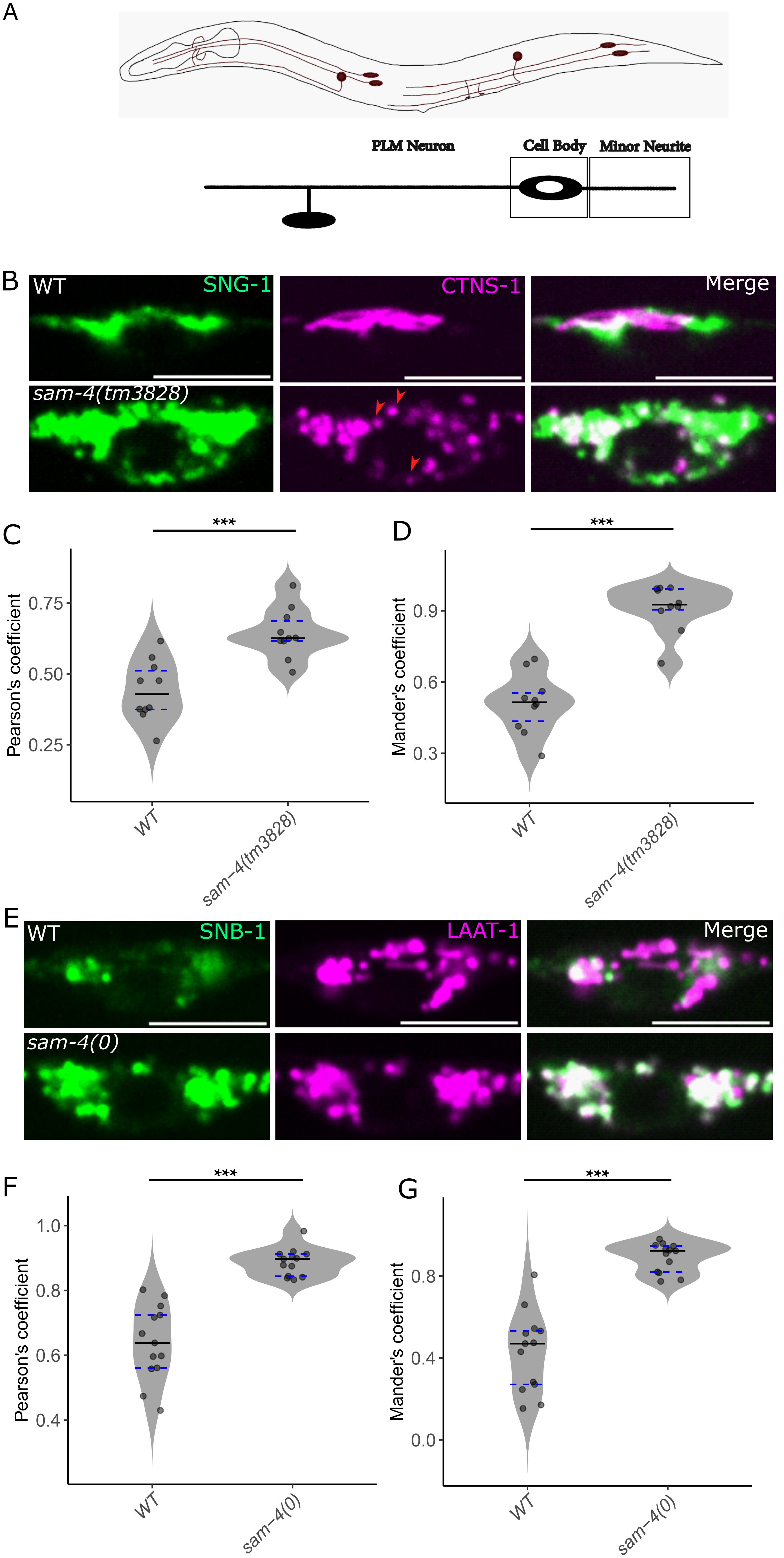

**Supplementary Figure 1**

A) Schematic representation of *C. elegans* showing the TRNs and the box indicates the region of imaging (PLM cell body and minor neurite).

B) Representative images of wild type and *sam-4(tm2388)* showing the overlap of SNG-1::mNeonGreen (Green) and CTNS-1::mCherry (Magenta) in the PLM cell body. Red arrowheads indicate fragmented CTNS-1 compartments. Scale bar = 5 µm.

C) Quantification of the overlap of SNG-1 and CTNS-1 by Pearson's Coefficient in wild type and *sam-4(tm3828)*. Number of animals ≥ 10, Statistical test : Unpaired Student t-test.

D) Quantification of the overlap of SNG-1 and CTNS-1 by Mander's Coefficient in wild type and *sam-4(tm3828)*. Number of animals ≥ 10, Statistical test : Wilcoxon rank-sum test.

E) Representative images of wild type and *sam-4(js415)* showing the overlap of SNB-1::eGFP (Green) and LAAT-1::mScarlet (Magenta) in the PLM cell body. Scale bar = 5 µm.

F) Quantification of the overlap of SNB-1 and LAAT-1 by Pearson's Coefficient in wild type and *sam-4(js415)*. Number of animals ≥ 12, Statistical test : Unpaired Student t-test.

G) Quantification of the overlap of SNB-1 and LAAT-1 by Mander's Coefficient in wild type and *sam-4(js415)*. Number of animals ≥ 12, Statistical test : Unpaired Student t-test.

All data plotted as violin plots showing individual data points marked along with the median (solid line), 25th and 75th percentile marked (dashed lines). * $p < 0.05$, ** $p < 0.01$; *** $p < 0.001$, ns - not significant.

### Supplementary Figure 2

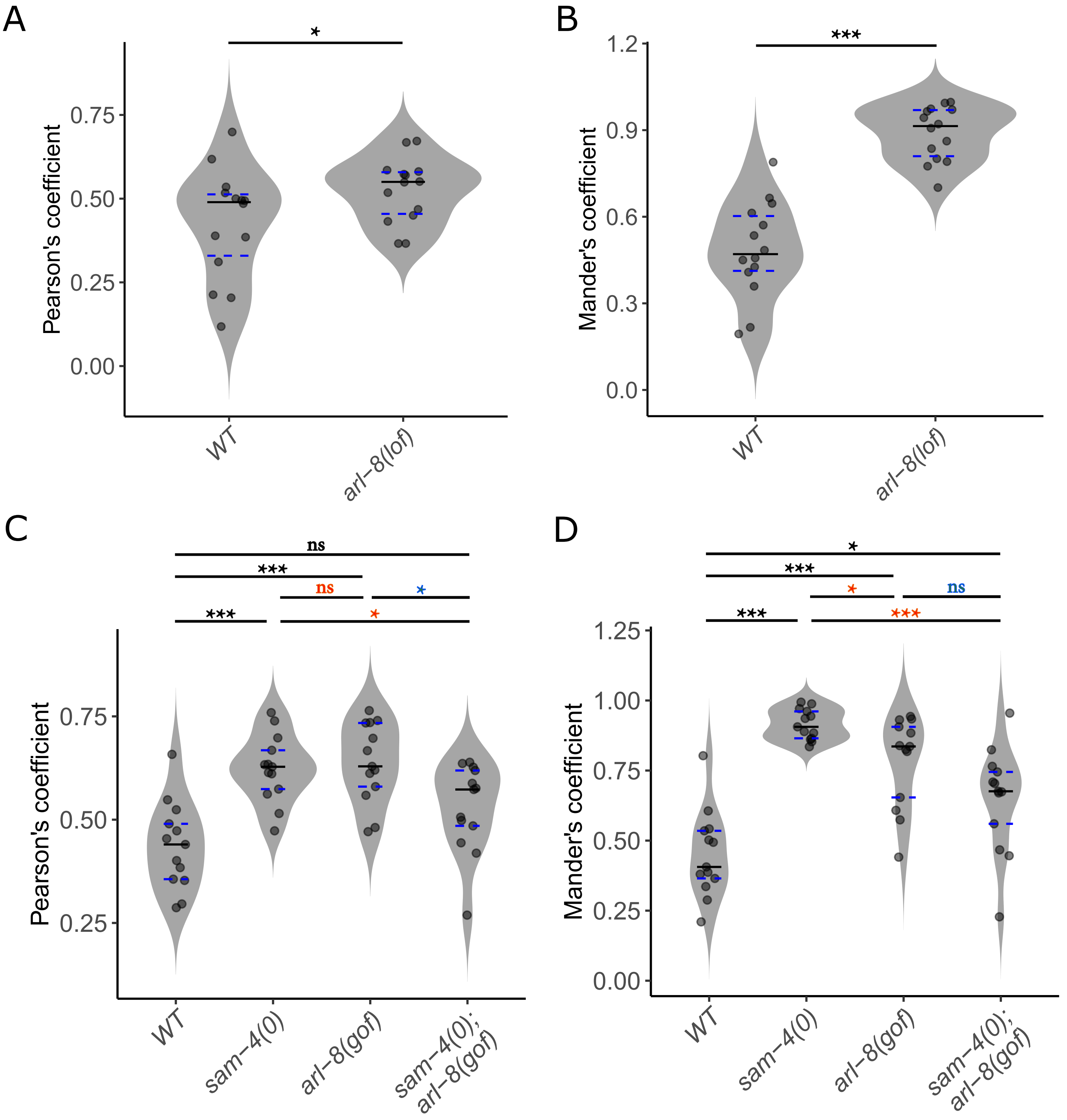

**Supplementary Figure 2**

A) Quantification of the overlap of SNG-1 and CTNS-1 by Pearson's Coefficient in wild type and *arl-8(tm2388)*. Number of animals ≥ 14, Statistical test : Unpaired Student t-test.

B) Quantification of the overlap of SNG-1 and CTNS-1 by Mander's Coefficient in wild type and *arl-8(tm2388)*. Number of animals ≥ 14, Statistical test : Wilcoxon rank-sum test.

C) Quantification of the overlap of SNG-1 and CTNS-1 by Pearson's Coefficient in wild type, *sam-4(js415), arl-8(jpn1), sam-4(js415); arl-8(jpn1)*. Number of animals ≥ 13, Statistical test : One way ANOVA, Post Hoc - Bonferroni correction.

D) Quantification of the overlap of SNG-1 and CTNS-1 by Mander's Coefficient in wild type, *sam-4(js415), arl-8(jpn1), sam-4(js415); arl-8(jpn1)*. Number of animals ≥ 13, One way ANOVA, Post Hoc - Bonferroni correction.

All data plotted as violin plots showing individual data points marked along with the median (solid line), 25th and 75th percentile marked (dashed lines). * p <0.05, ** p <0.01; *** p <0.001, ns - not significant. All relevant statistical comparisons are highlighted.

### Supplementary Figure 3

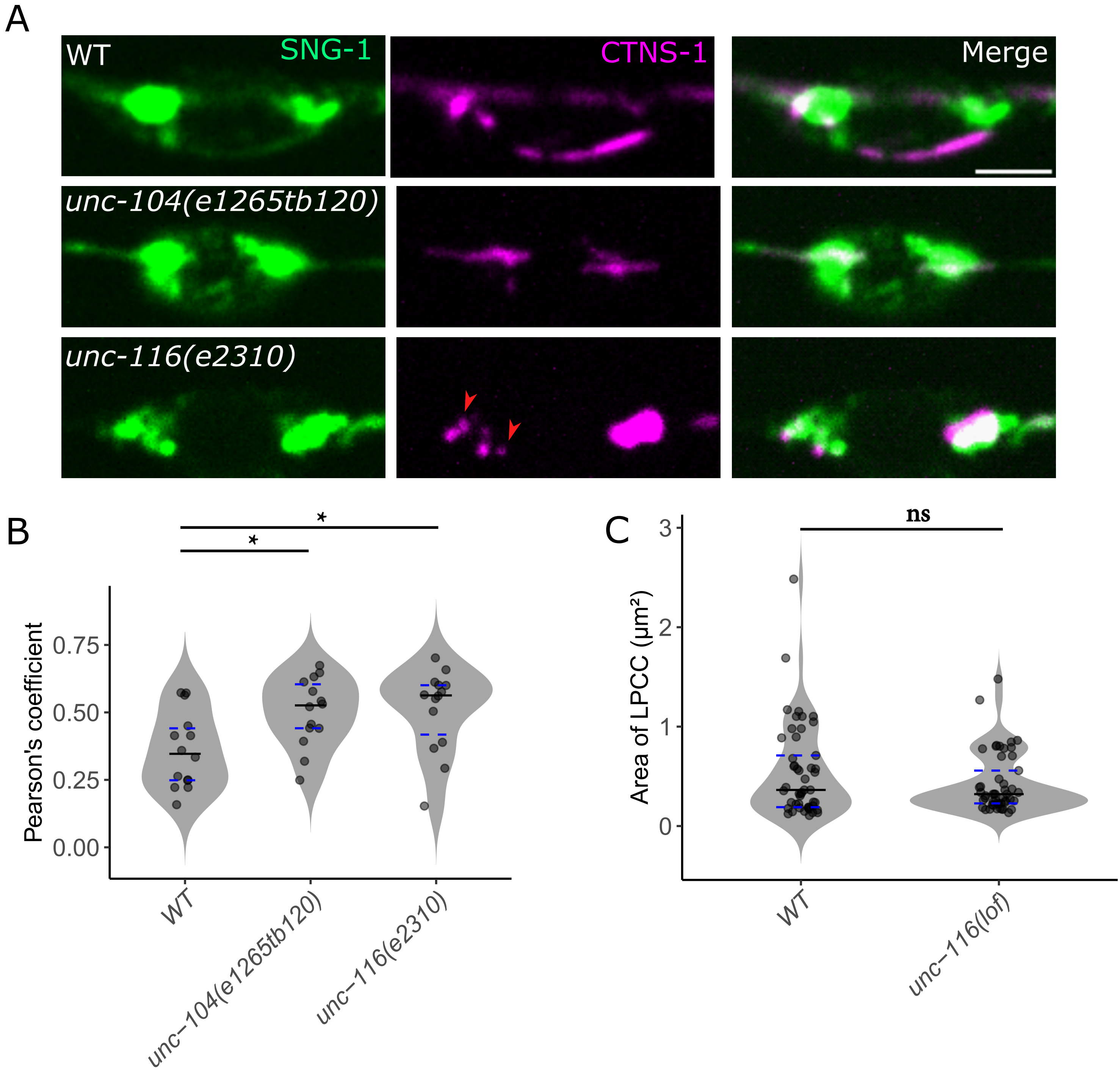

**Supplementary Figure 3**

A) Representative images of wild type, *unc-104(e1265tb120)*, and *unc-116(e2310)* showing the overlap of SNG-1::mNeonGreen (Green) and CTNS-1::mCherry (Magenta) in the PLM cell body. Red arrowheads indicate fragmented CTNS-1 compartments. Scale bar = 2.5 µm.
B) Quantification of the overlap of SNG-1 and CTNS-1 by Pearson's Coefficient in wild type, *unc-104(e1265tb120)*, and *unc-116(e2310)*. Number of animals ≥ 14, Statistical test : Wilcoxon rank-sum test.
C) Quantification of the area of dynamic CTNS-1 compartments in the minor neurite of PLM in wild type and *unc-104(e1265)*. Number of animals ≥ 11, Number of compartments ≥ 50, Statistical test : Kruskal Wallis ANOVA with Dunns test.

All data plotted as violin plots showing individual data points marked along with the median (solid line), 25th and 75th percentile marked (dashed lines). * $p < 0.05$, ** $p < 0.01$; *** $p < 0.001$, ns - not significant.

### Supplementary Figure 4

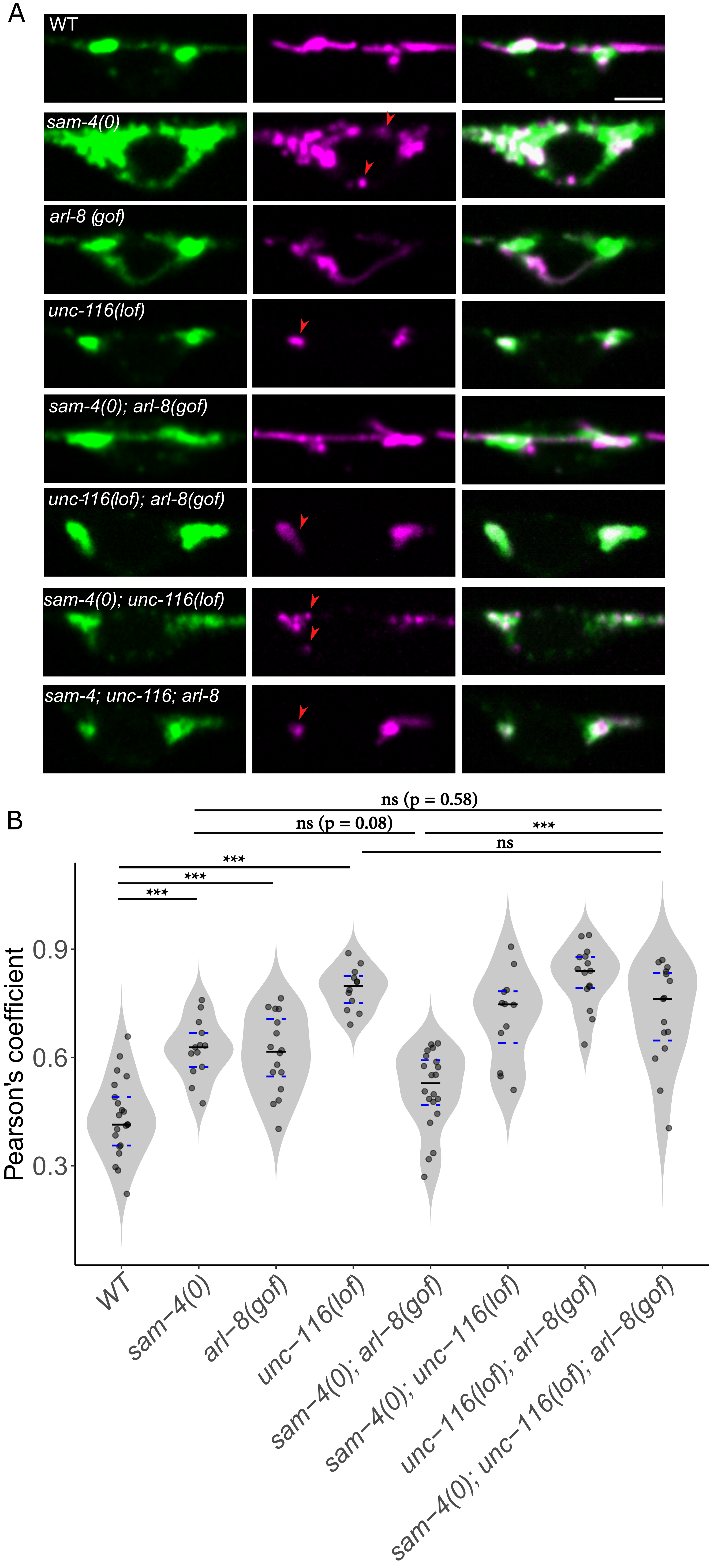

**Supplementary Figure 4**

A) Representative images of wild type, *sam-4(js415), arl-8(jpn1), unc-116(rh24sb79), sam-4(js415); arl-8(jpn1), unc-116(rh24sb79); arl-8(jpn1), sam-4(js415) unc-116(rh24sb79)* and *sam-4(js415); unc-116(rh24sb79); arl-8(jpn1)* showing the overlap of SNG-1::mNeonGreen (Green) and CTNS-1::mCherry (Magenta) in the PLM cell body. Red arrowheads indicate fragmented CTNS-1 compartments. Scale bar = 2.5 µm.

B) Quantification of the overlap of SNG-1 and CTNS-1 by Pearson's Coefficient in wild type, *sam-4(js415), arl-8(jpn1), unc-116(rh24sb79), sam-4(js415); arl-8(jpn1), unc-116(rh24sb79); arl-8(jpn1), sam-4(js415) unc-116(rh24sb79)* and *sam-4(js415); unc-116(rh24sb79); arl-8(jpn1)*. Number of animals $\geq$ 12, Statistical test : One way ANOVA, Post Hoc - Bonferroni correction.

All data plotted as violin plots showing individual data points marked along with the median (solid line), 25th and 75th percentile marked (dashed lines). * $p < 0.05$, ** $p < 0.01$; *** $p < 0.001$, ns - not significant. All relevant statistical comparisons are highlighted.

### Supplementary Figure 5

A

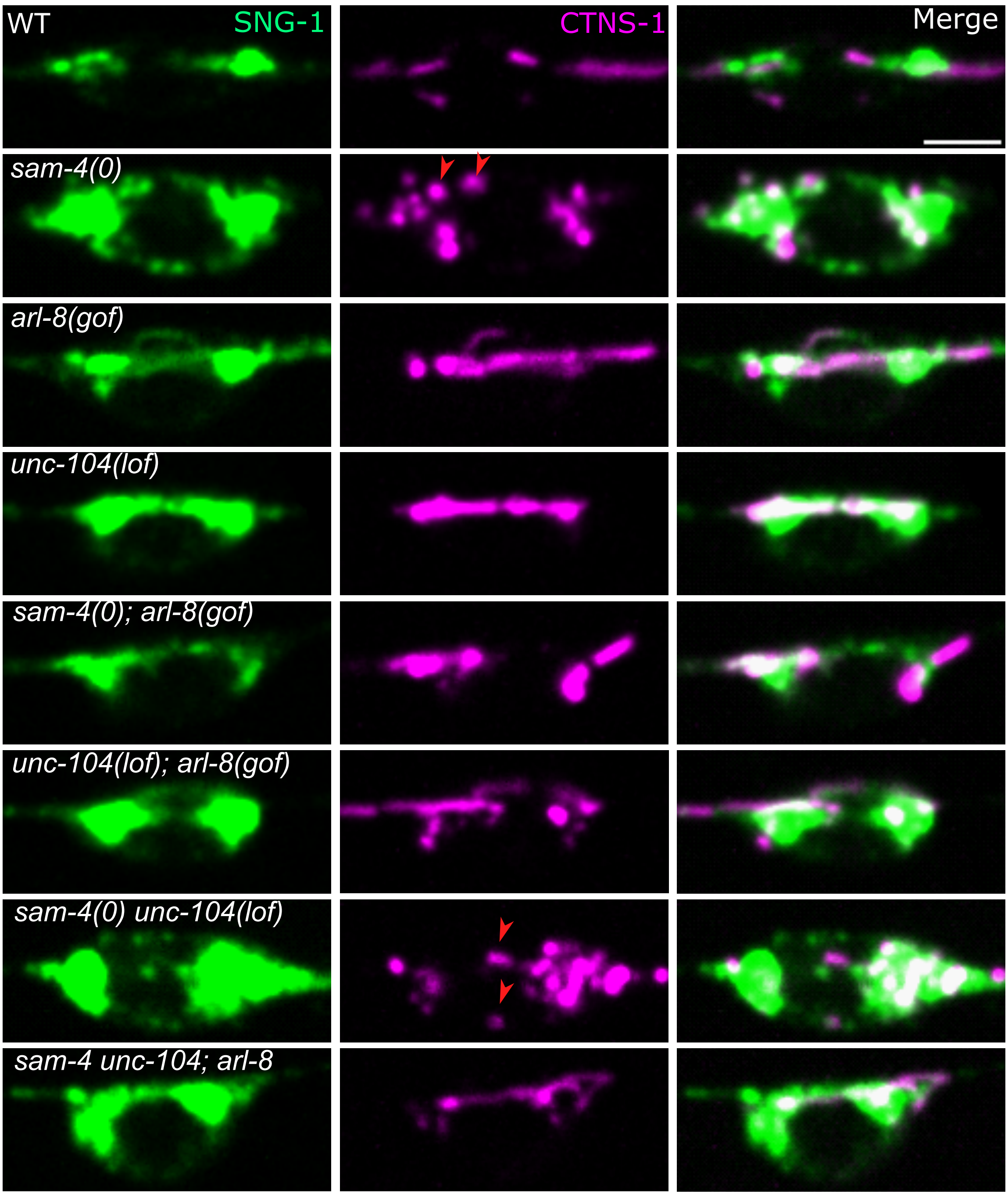


B

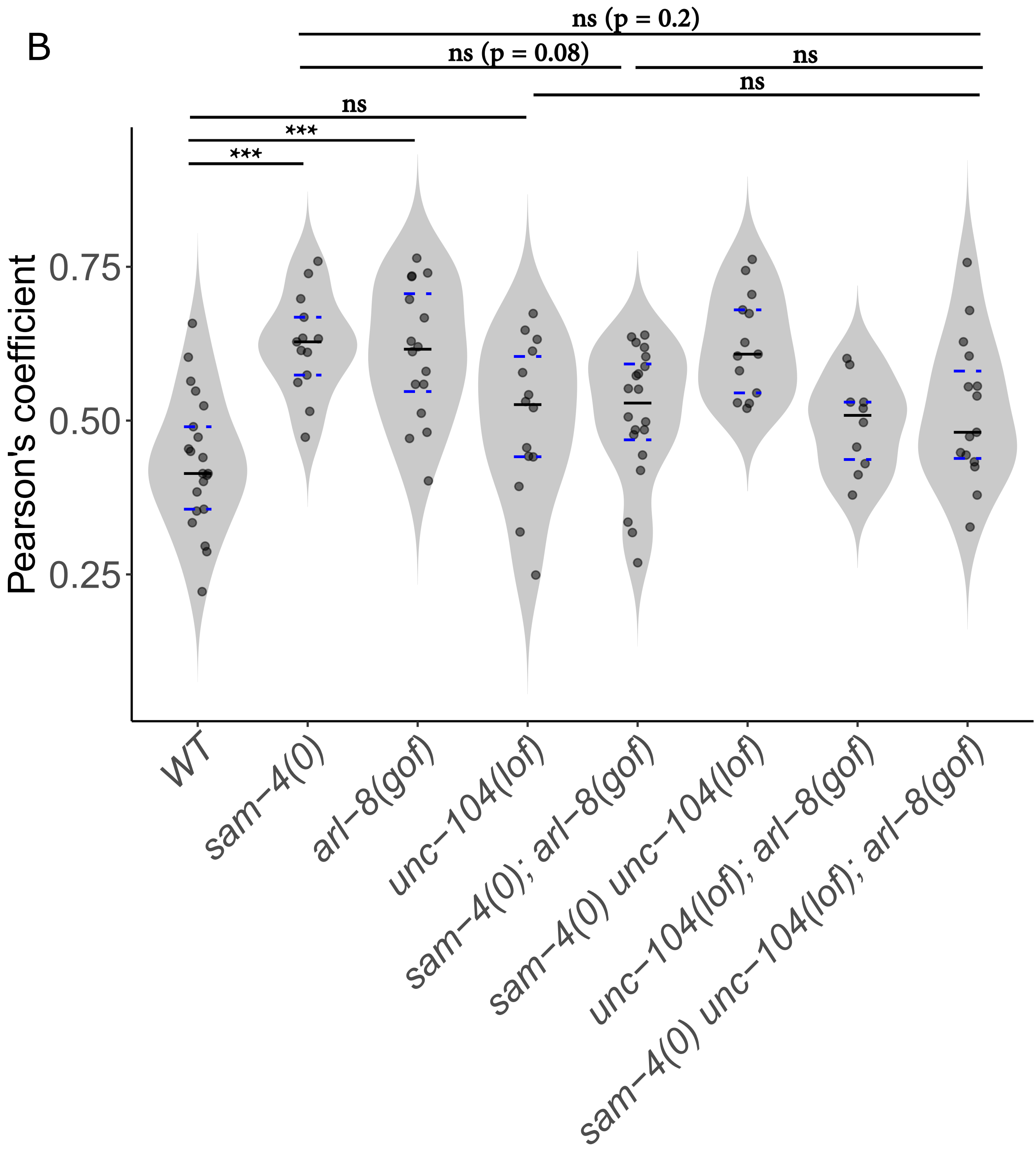

**Supplementary Figure 5**

A) Representative images of wild type, *sam-4(js415), arl-8(jpn1), unc-104(e1265tb120), sam-4(js415); arl-8(jpn1), unc-104(e1265tb120); arl-8(jpn1), sam-4(js415) unc-104(e1265tb120)* and *sam-4(js415) unc-104(e1265tb120); arl-8(jpn1)* showing the overlap of SNG-1::mNeonGreen (Green) and CTNS-1::mCherry (Magenta) in the PLM cell body. Red arrowheads indicate fragmented CTNS-1 compartments. Scale bar = 2.5 µm.

B) Quantification of the overlap of SNG-1 and CTNS-1 by Pearson's Coefficient in wild type, *sam-4(js415), arl-8(jpn1), unc-104(e1265tb120), sam-4(js415); arl-8(jpn1), unc-104(e1265tb120); arl-8(jpn1), sam-4(js415) unc-104(e1265tb120)* and *sam-4(js415) unc-104(e1265tb120); arl-8(jpn1)*. Number of animals ≥ 10, Statistical test : One way ANOVA, Post Hoc - Bonferroni correction.

All data plotted as violin plots showing individual data points marked along with the median (solid line), 25th and 75th percentile marked (dashed lines). * $p < 0.05$, ** $p < 0.01$; *** $p < 0.001$, ns - not significant. All relevant statistical comparisons are highlighted.

### Supplementary Figure 6

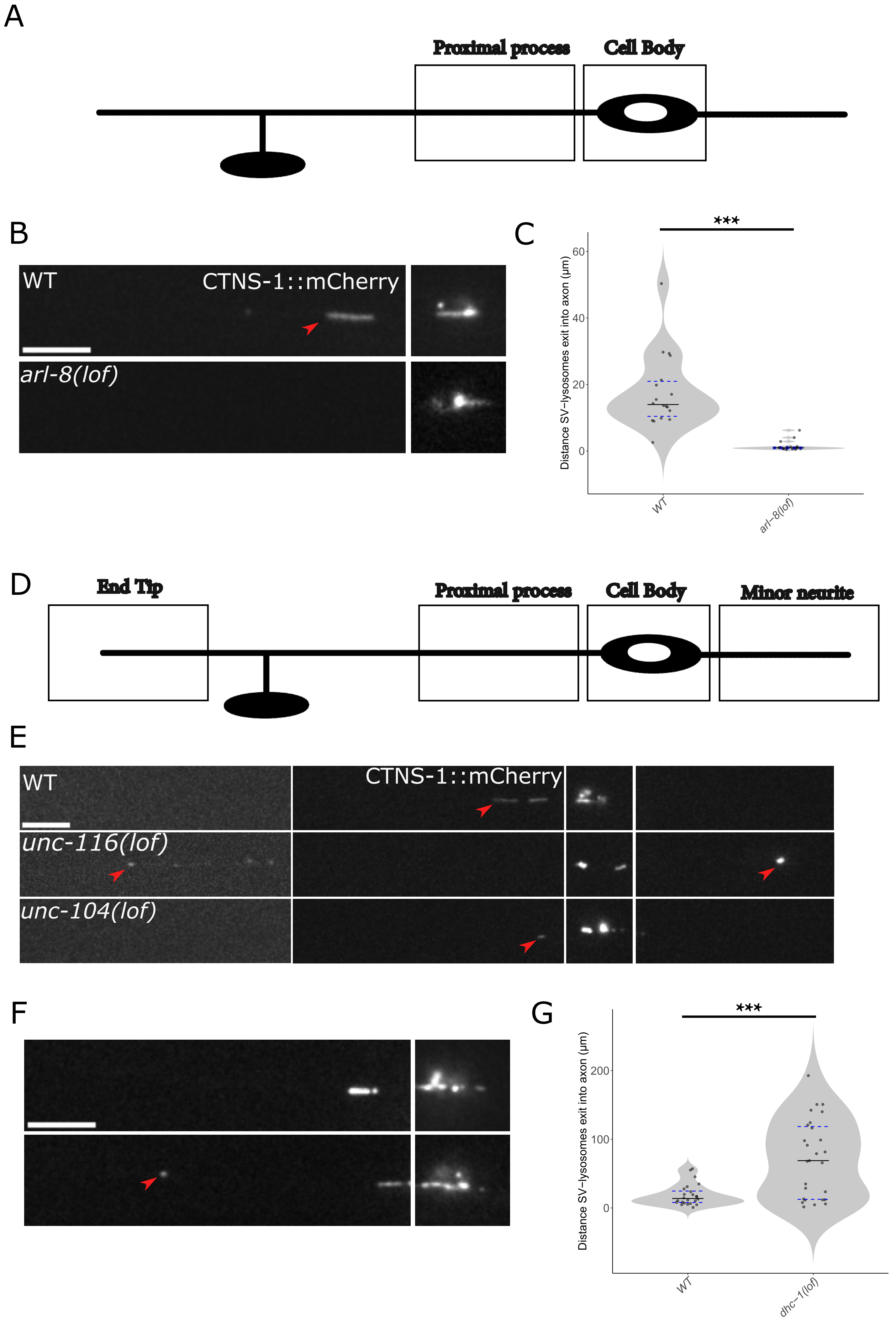

**Supplementary Figure 6**

A) Schematic representation of PLM and the box indicates the region of imaging.

B) Representative images of wild type and *arl-8(tm2388)* showing the distribution of CTNS-1::mCherry. Scale bar = 10 µm.

C) Quantification of the distance CTNS-1 exits into the neuronal process in wild type and *arl-8(tm2388)*. Number of animals ≥ 18, Statistical test : Wilcoxon rank-sum test.

D) Schematic representation of PLM and the box indicates the region of imaging.

E) Representative images of wild type, *unc-104(e1265tb120)* and *unc-116(rb24sb79)* showing the distribution of CTNS-1::mCherry. Scale bar = 10 µm.

F) Representative images of wild type and *dhc-1(js319)* showing the distribution of CTNS-1::mCherry. Scale bar = 10 µm.

G) Quantification of the distance CTNS-1 exits into the neuronal process in wild type and *dhc-1(js319)*. Number of animals ≥ 20, Statistical test : Wilcoxon rank-sum test.

All data plotted as violin plots showing individual data points marked along with the median (solid line), 25th and 75th percentile marked (dashed lines). * p <0.05, ** p <0.01; *** p <0.001, ns - not significant.

### Supplementary Figure 7

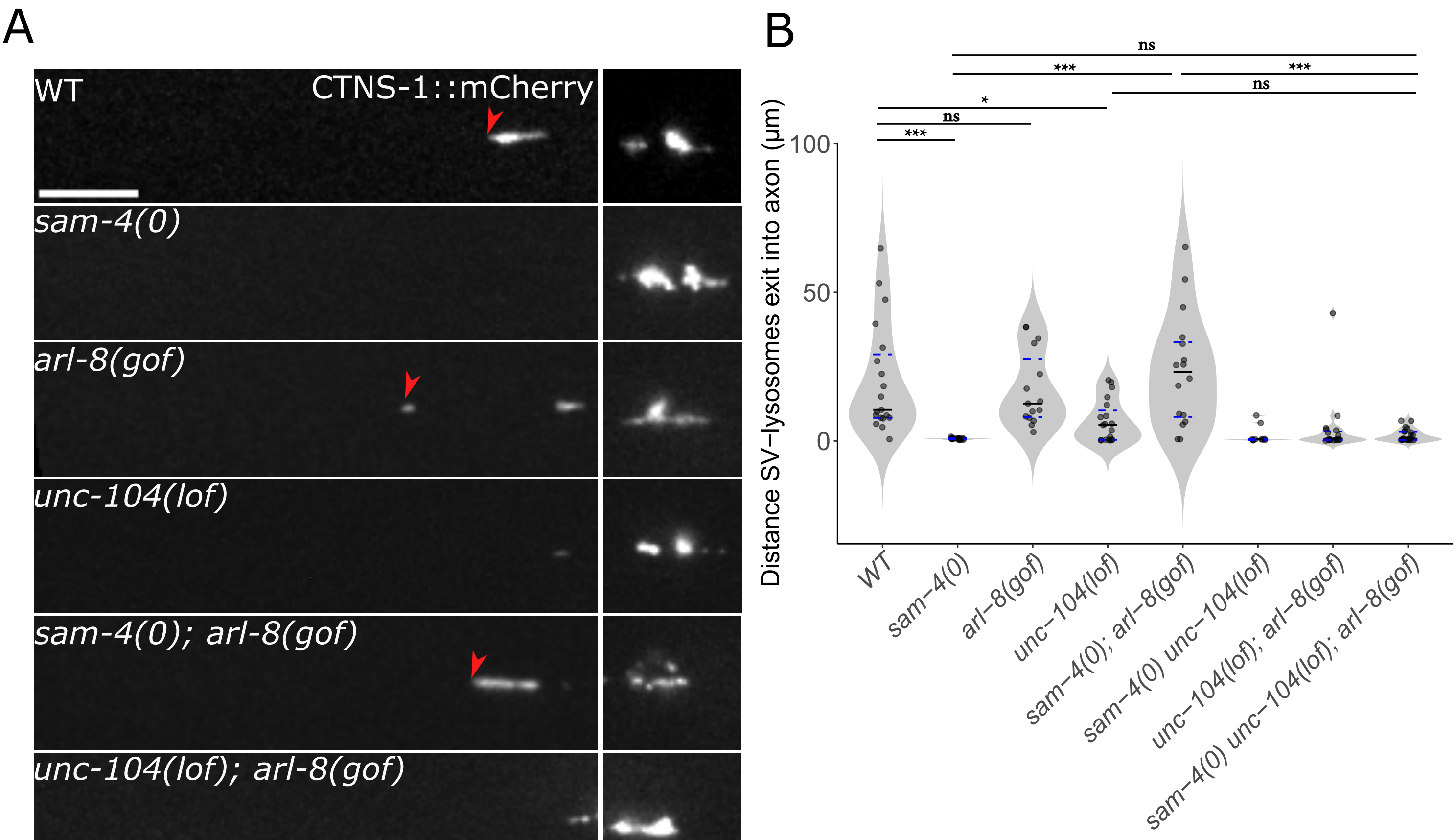

**Supplementary Figure 7**

A) Representative images of wild type, *sam-4(js415), arl-8(jpn1), unc-104(e1265tb120), sam-4(js415); arl-8(jpn1), unc-104(e1265tb120); arl-8(jpn1), sam-4(js415) unc-104(e1265tb120)* and *sam-4(js415) unc-104(e1265tb120); arl-8(jpn1)* showing the distribution of CTNS-1::mCherry. Red arrowhead indicates the distance CTNS-1 exits into the neuronal process. Scale bar = 10 µm.

B) Quantification of the distance CTNS-1 exits into the neuronal process in wild type, *sam-4(js415), arl-8(jpn1), unc-104(e1265tb120), sam-4(js415); arl-8(jpn1), unc-104(e1265tb120); arl-8(jpn1), sam-4(js415) unc-104(e1265tb120)* and *sam-4(js415) unc-104(e1265tb120); arl-8(jpn1)*. Number of animals ≥ 11, Statistical test : Kruskal Wallis ANOVA with Dunns test.

All data plotted as violin plots showing individual data points marked along with the median (solid line), 25th and 75th percentile marked (dashed lines). * p <0.05, ** p <0.01; *** p <0.001, ns - not significant.

### Supplementary Figure 8

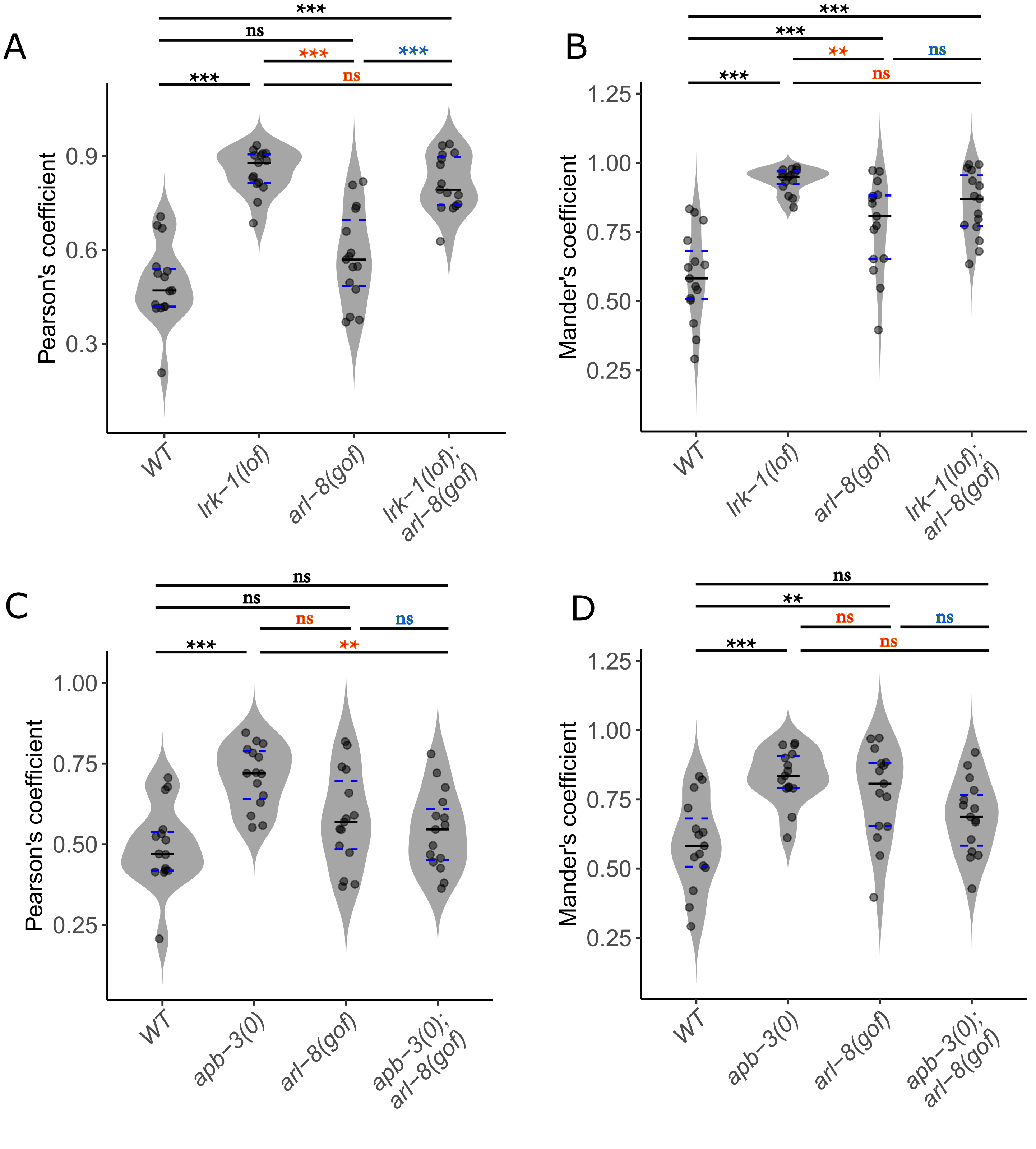

**Supplementary Figure 8**

A) Quantification of the overlap of SNG-1 and CTNS-1 by Pearson's Coefficient in wild type, *lrk-1(km17), arl-8(jpn1)* and *lrk-1(km17); arl-8(jpn1)*. Number of animals ≥ 15, Statistical test : One way ANOVA, Post Hoc - Bonferroni correction.

B) Quantification of the overlap of SNG-1 and CTNS-1 by Mander's Coefficient in wild type, *lrk-1(km17), arl-8(jpn1)* and *lrk-1(km17); arl-8(jpn1)*. Number of animals ≥ 15, Statistical test : One way ANOVA, Post Hoc - Bonferroni correction.

C) Quantification of the overlap of SNG-1 and CTNS-1 by Pearson's Coefficient in wild type, *apb-3(ok429), arl-8(jpn1)* and *apb-3(ok429); arl-8(jpn1)*. Number of animals ≥ 15, Statistical test : One way ANOVA, Post Hoc - Bonferroni correction.

D) Quantification of the overlap of SNG-1 and CTNS-1 by Mander's Coefficient in wild type, *apb-3(ok429), arl-8(jpn1)* and *apb-3(ok429); arl-8(jpn1)*. Number of animals ≥ 15, Statistical test : One way ANOVA, Post Hoc - Bonferroni correction.

All data plotted as violin plots showing individual data points marked along with the median (solid line), 25th and 75th percentile marked (dashed lines). * p <0.05, ** p <0.01; *** p <0.001, ns - not significant. All relevant statistical comparisons are highlighted.

### Supplementary Figure 9

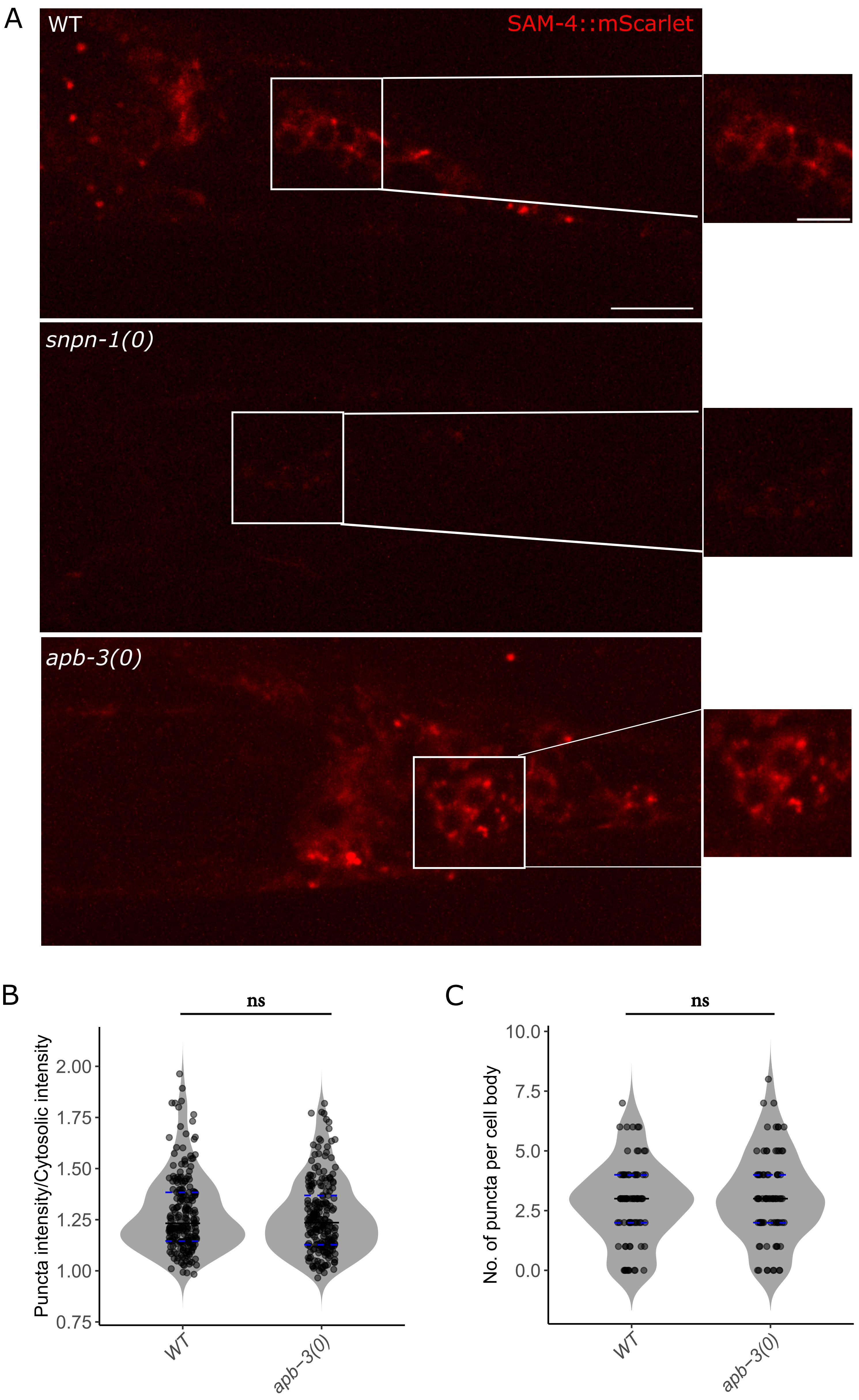

**Supplementary Figure 9**

A) Representative images of wild type, *snpn-1(tm1892) and apb-3(ok429)* showing the localisation of SAM-4::mScarlet in the cell bodies at the tail of *C. elegans*. Scale bar =10 µm (zoomed out panel), 2.5 µm (zoomed in panel).

B) Quantification of the intensity (Puncta intensity/Cytosolic intensity) of SAM-4 puncta from the tail cell bodies in wild type and *apb-3(ok429)*. Number of animals ≥ 10, Number of cell bodies ≥ 81, Number of puncta ≥ 228, Statistical test : Wilcoxon rank-sum test.

C) Quantification of the number of SAM-4 puncta per cell body from the tail cell bodies in wild type and *apb-3(ok429)*. Number of animals ≥ 10, Number of cell bodies ≥ 81, Statistical test : Wilcoxon rank-sum test.

All data plotted as violin plots showing individual data points marked along with the median (solid line), 25th and 75th percentile marked (dashed lines). * $p < 0.05$, ** $p < 0.01$; *** $p < 0.001$, ns - not significant. All relevant statistical comparisons are highlighted.

### Supplementary Figure 10

A Wild type

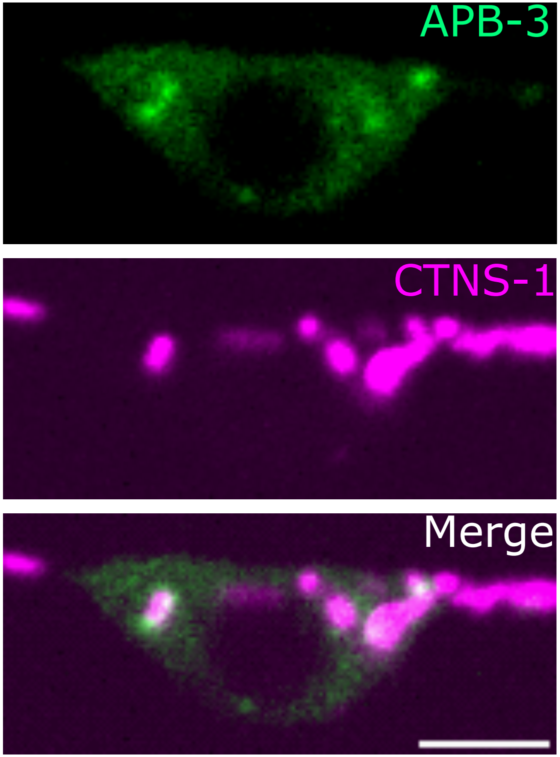


B

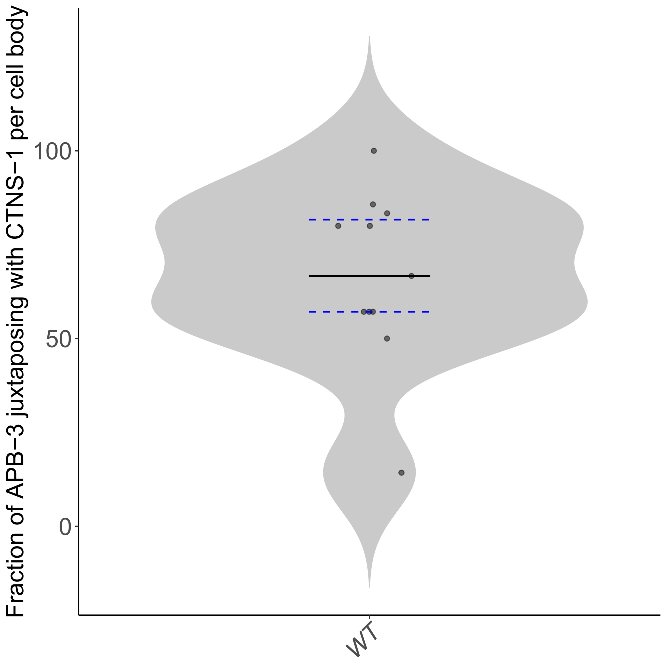

**Supplementary Figure 10**

A) Representative images showing the juxtaposition of APB-3::GFP (Green) and CTNS-1::mCherry (Magenta) in wild type animals. Scale bar = 2.5 μm.
B) Quantification of the juxtaposition of APB-3::GFP and CTNS-1::mCherry in wild type animals. Number of animals =11.

All data plotted as violin plots showing individual data points marked along with the median (solid line), 25th and 75th percentile marked (dashed lines).
