## Supplementary Table 1 for "BORC segregates synaptic vesicle and lysosomal proteins through motors UNC-104/KIF1A and UNC-116/KIF5"

**Table S1**

| Sl.No | Strain number | Genotype | Reference |
|---|---|---|---|
| 1 | N2 | Bristol wild type | [1] |
| 2 | TT994 | *sam-4(js415)* II null *(0)* | [2] |
| 3 | TT1045 | *sam-4(tm3828)* II null *(0)* | [3] |
| 4 | TT2424 | *dsbn-1 (tm6580)* I loss of function *(lof)* | [3] |
| 5 | TT2425 | *snpn-1(tm1892)* IV null *(0)* | [3] |
| 6 | TT1482 | *arl-8(tm2388)* IV loss of function *(lof)* | [3] |
| 7 | TT740 | *arl-8(jpn1)* IV gain of function *(gof)* | [4] |
| 8 | TT397 | *unc-104(e1265)* II loss of function *(lof)* | [5] |
| 9 | TT385 | *unc-104(e1265tb120)* II loss of function *(lof)* | [6] |
| 10 | TT174 | *unc-116(e2310)* III loss of function *(lof)* | [7] |
| 11 | TT189 | *unc-116(rh24sb79)* III loss of function *(lof)* | [8] |
| 12 | TT388 | *dhc-1(js319)* I loss of function *(lof)* | [9] |
| 13 | TT2060 | *apb-3(ok429)* I null *(0)* | [10] |
| 14 | TT1218 | *lrk-1 (km17)* I loss of function *(lof)* | [11] |
| 15 | TT2903 | *tbIs381* [*mec-4*p::*ctns-1*::*mCherry (20 ng/µl), myo-2p::H2b::GFP (40 ng/µl), pBluescript SK (110 ng/µl)]* X | [12] |
| 16 | TT2884 | *tbIs388* [*mec-4*p::*sng-1*::*gfp (5 ng/µl), myo-2p::mCherry (10 ng/µl), pBluescript SK (185 ng/µl)]* | [12] |
| 17 | TT3477 | *tbSi520 [mec-4S*p *sng-1-L-mNG]* IV (Single copy) | [13] |
| 18 | TT3007 | *tbIs414* [*mec-7*p::*snb-1*::*egfp (10 ng/µl), myo-2p::mCherry (10 ng/µl), pBluescript SK (180 ng/µl)]* | [12] |
| 19 | TT3586 | *jsSi2052 [mec-7b*p *laat-1-L-Scarlet]* V (Single copy) | This study |
| 20 | TT3608 | *tbIs518 [rab-3p::apb-3::gfp (30 ng/µl), myo-2p::mCherry (10 ng/µl), pBluescript SK (140 ng/µl)]* | [12] |
| 21 | WEH644 | *sam-4(syb4105*[*sam-4::mScarlet-I::ZF1*]) II (CRISPR) | [14] |
