## Supplementary material for "BORC segregates synaptic vesicle and lysosomal proteins through motors UNC-104/KIF1A and UNC-116/KIF5": Movie legends

**Supplementary movie legends**

**Movie 1 : Z stacks of SNG-1::mNeonGreen and CTNS-1::mCherry in wild type animals**

Z stacks of SNG-1::mNeonGreen (Green) and CTNS-1::mCherry (Magenta) in the PLM cell body of wild type animals. Z slice interval of 0.28um and playback speed 4 slices per second. Genotype: *tbSi520; tbIs381,* Scale: 2.5 µm.

**Movie 2 : Z stacks of SNG-1::mNeonGreen and CTNS-1::mCherry in *sam-4(0)* animals**

Z stacks of SNG-1::mNeonGreen (Green) and CTNS-1::mCherry (Magenta) in the PLM cell body of *sam-4(js415)* animals. Z slice interval of 0.28um and playback speed 4 slices per second. Genotype: *sam-4(js415); tbSi520; tbIs381,* Scale: 2.5 µm.

**Movie 3 : Dynamic SNG-1::GFP and CTNS-1::mCherry in wild type animals**

Dynamic SNG-1::GFP (Green) and CTNS-1::mCherry (Magenta) in the PLM cell body of wild type animals. Imaged sequentially at 2.3 frames per second (fps), playback speed 20 fps. Genotype: *tbIs388; tbIs381,* Scale: 2.5 µm.

**Movie 4 : Dynamic SNG-1::GFP and CTNS-1::mCherry in *snpn-1(0)* animals**

Dynamic SNG-1::GFP(Green) and CTNS-1::mCherry (Magenta) in the PLM cell body of wild type animals. Imaged sequentially at 2.3 frames per second (fps), playback speed 20 fps. Genotype: *snpn-1; tbIs388; tbIs381,* Scale: 2.5 µm.

**Movie 5 : Z stacks of SNB-1::GFP and LAAT-1::mScarlet in wild type animals**

Z stacks of SNB-1::GFP (Green) and LAAT-1::mScarlet (Magenta) in the PLM cell body of wild type animals. Z slice interval of 0.28um and playback speed 4 slices per second. Genotype: *tbIs414; jsSi2052,* Scale: 2.5 µm.

**Movie 6 : Z stacks of SNB-1::GFP and LAAT-1::mScarlet in *sam-4(0)* animals**

Z stacks of SNB-1::GFP (Green) and LAAT-1::mScarlet (Magenta) in the PLM cell body of *sam-4(js415)* animals. Z slice interval of 0.28um and playback speed 4 slices per second. Genotype: *sam-4(js415); tbIs414; jsSi2052,* Scale: 2.5 µm.

**Movie 7 : Dynamic SNG-1::mNeonGreen and CTNS-1::mCherry in wild type animals**

Dynamic SNG-1::mNeonGreen (Green) and CTNS-1::mCherry (Magenta) in the PLM cell body of wild type animals. Imaged sequentially at 2.3 frames per second (fps), playback speed 20 fps. Genotype: *tbSi520; tbIs381,* Scale: 2.5 µm.

**Movie 8 : Dynamic SNG-1::mNeonGreen and CTNS-1::mCherry in *sam-4(0)* animals**

Dynamic SNG-1::mNeonGreen (Green) and CTNS-1::mCherry (Magenta) in the PLM cell body of *sam-4(js415)* animals. Imaged sequentially at 2.3 frames per second (fps), playback speed 20 fps. Genotype: *sam-4(js415); tbSi520; tbIs381,* Scale: 2.5 µm.

**Movie 9 : Z stacks of SNG-1::mNeonGreen and CTNS-1::mCherry in *arl-8(lof)* animals**

Z stacks of SNG-1::mNeonGreen (Green) and CTNS-1::mCherry (Magenta) in the PLM cell body of *arl-8(tm2388)* animals. Z slice interval of 0.28um and playback speed 4 slices per second. Genotype: *arl-8(tm2388); tbSi520; tbIs381,* Scale: 2.5 µm.

**Movie 10 : Dynamic SNG-1::mNeonGreen and CTNS-1::mCherry in *arl-8(lof)* animals**

Dynamic SNG-1::mNeonGreen (Green) and CTNS-1::mCherry (Magenta) in the PLM cell body of *arl-8(tm2388)* animals. Imaged sequentially at 2.3 frames per second (fps), playback speed 20 fps. Genotype: *arl-8(tm2388); tbSi520; tbIs381,* Scale: 2.5 µm.

**Movie 11 : Z stacks of SNG-1::mNeonGreen and CTNS-1::mCherry in *arl-8(gof)* animals**

Z stacks of SNG-1::mNeonGreen (Green) and CTNS-1::mCherry (Magenta) in the PLM cell body of *arl-8(jpn1)* animals. Z slice interval of 0.28um and playback speed 4 slices per second. Genotype: *arl-8(jpn1); tbSi520; tbIs381,* Scale: 2.5 µm.

**Movie 12 : Z stacks of SNG-1::mNeonGreen and CTNS-1::mCherry in *sam-4(0); arl-8(gof)* animals**

Z stacks of SNG-1::mNeonGreen (Green) and CTNS-1::mCherry (Magenta) in the PLM cell body of *sam-4(js415); arl-8(jpn1)* animals. Z slice interval of 0.28um and playback speed 4 slices per second. Genotype: *sam-4(js415); arl-8(jpn1); tbSi520; tbIs381,* Scale: 2.5 µm.

**Movie 13 : Dynamic SNG-1::mNeonGreen and CTNS-1::mCherry in *arl-8(gof)* animals**

Dynamic SNG-1::mNeonGreen (Green) and CTNS-1::mCherry (Magenta) in the PLM cell body of *arl-8(jpn1)* animals. Imaged sequentially at 2.3 frames per second (fps), playback speed 20 fps. Genotype: *arl-8(jpn1); tbSi520; tbIs381,* Scale: 2.5 µm.

**Movie 14 : Dynamic SNG-1::mNeonGreen and CTNS-1::mCherry in *sam-4(0); arl-8(gof)* animals**

Dynamic SNG-1::mNeonGreen (Green) and CTNS-1::mCherry (Magenta) in the PLM cell body of *sam-4(js415); arl-8(jpn1)* animals. Imaged sequentially at 2.3 frames per second (fps), playback speed 20 fps. Genotype: *sam-4(js415); arl-8(jpn1); tbSi520; tbIs381,* Scale: 2.5 µm.

**Movie 15 : Z stacks of SNG-1::mNeonGreen and CTNS-1::mCherry in *unc-104(lof)* animals**

Z stacks of SNG-1::mNeonGreen (Green) and CTNS-1::mCherry (Magenta) in the PLM cell body of *unc-104(e1265)* animals. Z slice interval of 0.28um and playback speed 4 slices per second. Genotype: *unc-104(e1265); tbSi520; tbIs381,* Scale: 2.5 µm.

**Movie 16 : Z stacks of SNG-1::mNeonGreen and CTNS-1::mCherry in *unc-116(lof)* animals**

Z stacks of SNG-1::mNeonGreen (Green) and CTNS-1::mCherry (Magenta) in the PLM cell body of *unc-116(rh24sb79)* animals. Z slice interval of 0.28um and playback speed 4 slices per second. Genotype: *unc-116(rh24sb79); tbSi520; tbIs381,* Scale: 2.5 µm.

**Movie 17 : Dynamic SNG-1::mNeonGreen and CTNS-1::mCherry in *unc-104(lof)* animals**

Dynamic SNG-1::mNeonGreen (Green) and CTNS-1::mCherry (Magenta) in the PLM cell body of *unc-104(e1265)* animals. Imaged sequentially at 2.3 frames per second (fps), playback speed 20 fps. Genotype: *unc-104(e1265); tbSi520; tbIs381,* Scale: 2.5 µm.

**Movie 18 : Dynamic SNG-1::mNeonGreen and CTNS-1::mCherry in *unc-116(lof)* animals**

Dynamic SNG-1::mNeonGreen (Green) and CTNS-1::mCherry (Magenta) in the PLM cell body of *unc-116(rh24sb79)* animals. Imaged sequentially at 2.3 frames per second (fps), playback speed 20 fps. Genotype: *unc-116(rh24sb79); tbSi520; tbIs381,* Scale: 2.5 µm.

**Movie 19 : Dynamic SNG-1::mNeonGreen and CTNS-1::mCherry in the minor neurite of wild type animals**

Dynamic SNG-1::mNeonGreen (Green) and CTNS-1::mCherry (Magenta) in the minor neurite of PLM in wild type animals. Imaged sequentially at 2.3 frames per second (fps), playback speed 20 fps. Genotype: *tbSi520; tbIs381,* Scale: 10 µm.

**Movie 20 : Dynamic SNG-1::mNeonGreen and CTNS-1::mCherry in the minor neurite of *unc-116(lof)* animals**

Dynamic SNG-1::mNeonGreen (Green) and CTNS-1::mCherry (Magenta) in the minor neurite of PLM in *unc-116(rh24sb79)* animals. Imaged sequentially at 2.3 frames per second (fps), playback speed 20 fps. Genotype: *unc-116(rh24sb79); tbSi520; tbIs381,* Scale: 10 µm.

**Movie 21 : Z stacks of SNG-1::mNeonGreen and CTNS-1::mCherry in *lrk-1(lof)* animals**

Z stacks of SNG-1::mNeonGreen (Green) and CTNS-1::mCherry (Magenta) in the PLM cell body of *lrk-1(km17)* animals. Z slice interval of 0.28um and playback speed 4 slices per second. Genotype: *lrk-1(km17); tbSi520; tbIs381,* Scale: 2.5 µm.

**Movie 22 : Dynamic SNG-1::mNeonGreen and CTNS-1::mCherry in *lrk-1(lof)* animals**

Dynamic SNG-1::mNeonGreen (Green) and CTNS-1::mCherry (Magenta) in the PLM cell body of *lrk-1(km17)* animals. Imaged sequentially at 2.3 frames per second (fps), playback speed 20 fps. Genotype: *lrk-1(km17); tbSi520; tbIs381,* Scale: 2.5 µm.

**Movie 23 : Dynamic SNG-1::mNeonGreen and CTNS-1::mCherry in *lrk-1(lof); arl-8(gof)* animals**

Dynamic SNG-1::mNeonGreen (Green) and CTNS-1::mCherry (Magenta) in the PLM cell body of *lrk-1(km17); arl-8(jpn1)* animals. Imaged sequentially at 2.3 frames per second (fps), playback speed 20 fps. Genotype: *lrk-1(km17); arl-8(jpn1); tbSi520; tbIs381,* Scale: 2.5 µm.

**Movie 24 : Z stacks of SNG-1::mNeonGreen and CTNS-1::mCherry in *lrk-1(lof); arl-8(gof)* animals**

Z stacks of SNG-1::mNeonGreen (Green) and CTNS-1::mCherry (Magenta) in the PLM cell body of *lrk-1(km17); arl-8(jpn1)* animals. Z slice interval of 0.28um and playback speed 4 slices per second. Genotype: *lrk-1(km17); arl-8(jpn1); tbSi520; tbIs381,* Scale: 2.5 µm.

**Movie 25 : Z stacks of SNG-1::mNeonGreen and CTNS-1::mCherry in *apb-3(lof)* animals**

Z stacks of SNG-1::mNeonGreen (Green) and CTNS-1::mCherry (Magenta) in the PLM cell body of *apb-3(ok429)* animals. Z slice interval of 0.28um and playback speed 4 slices per second. Genotype: *apb-3(ok429); tbSi520; tbIs381,* Scale: 2.5 µm.

**Movie 26 : Dynamic SNG-1::mNeonGreen and CTNS-1::mCherry in *apb-3(lof)* animals**

Dynamic SNG-1::mNeonGreen (Green) and CTNS-1::mCherry (Magenta) in the PLM cell body of *apb-3(ok429)* animals. Imaged sequentially at 2.3 frames per second (fps), playback speed 20 fps. Genotype: *apb-3(ok429); tbSi520; tbIs381,* Scale: 2.5 µm.

**Movie 27 : Dynamic SNG-1::mNeonGreen and CTNS-1::mCherry in *apb-3(lof); arl-8(gof)* animals**

Dynamic SNG-1::mNeonGreen (Green) and CTNS-1::mCherry (Magenta) in the PLM cell body of *apb-3(ok429); arl-8(jpn1)* animals. Imaged sequentially at 2.3 frames per second (fps), playback speed 20 fps. Genotype: *apb-3(ok429); arl-8(jpn1); tbSi520; tbIs381,* Scale: 2.5 µm.

**Movie 28 : Z stacks of SNG-1::mNeonGreen and CTNS-1::mCherry in *apb-3(lof); arl-8(gof)* animals**

Z stacks of SNG-1::mNeonGreen (Green) and CTNS-1::mCherry (Magenta) in the PLM cell body of *apb-3(ok429); arl-8(jpn1)* animals. Z slice interval of 0.28um and playback speed 4 slices per second. Genotype: *apb-3(ok429); arl-8(jpn1); tbSi520; tbIs381,* Scale: 2.5 µm.

**Movie 29 : Z stacks of APB-3::GFP and CTNS-1::mCherry in wild type animals**

Z stacks of APB-3::GFP (Green) and CTNS-1::mCherry (Magenta) in the PLM cell body of wild type animals. Z slice interval of 0.28um and playback speed 4 slices per second. Genotype: *tbIs518; tbIs381*, Scale: 2.5 µm.
